# Purification of post-transcriptionally modified tRNAs for enhanced cell-free translation systems

**DOI:** 10.1101/2025.06.10.658963

**Authors:** Evan M. Kalb, Jose L. Alejo, Leticia Dias-Fields, Isaac Knudson, Joshua A. Davisson, Efren Maldonado, Kanokporn Chattrakun, Shangsi Lin, Alanna Schepartz, Shenglong Zhang, Scott Blanchard, Aaron E. Engelhart, Katarzyna P. Adamala

**Affiliations:** Department of Genetics, Cell Biology and Development, University of Minnesota, Minneapolis, MN US; Department of Chemistry, University of California, Berkeley, CA, USA; Department of Structural Biology, St. Jude Children’s Research Hospital, Memphis, Tennessee, USA; NSF Center for Genetically Encoded Materials (C-GEM), Berkeley, CA, USA; Department of Chemistry and The RNA Institute, University at Albany, State University of New York, Albany, NY 12222, USA

## Abstract

Transfer RNAs (tRNAs) are utilized by the ribosome to decode the nucleic acid alphabet. tRNA structure, stability, aminoacylation efficiency, and decoding efficacy are governed by their extensive post-transcriptional modifications. In most studies, individual tRNAs are generated using *in vitro* transcription, which produces tRNAs devoid of these critical site-specific modifications, negatively affecting translation yields and fidelity. To address this, we have developed a purification method which couples tRNA overexpression to DNA hybridization-based purification. Using this approach, we produced native tRNAs from *E. coli* in high yield and purity while retaining their complement of native post-transcriptional modifications and translational activity. We extend this technique to the purification of 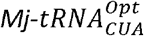 and 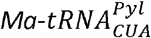, tRNAs of critical importance for genetic code expansion. We confirmed that both 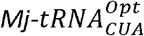 and 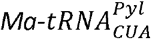 contain native *E. coli* post-transcriptional modifications and provide the first complete modification profiles of each. Moreover, we found that *in vivo-*generated 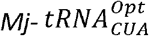 significantly outperforms its *in vitro-*generated counterpart in amber codon suppression in cell-free translation reactions. Finally, we purified an engineered variant of *E. coli* 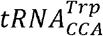, extending our studies to synthetic tRNAs. We present a flexible method which generates modified tRNAs in high yield and purity, addressing a critical and persistent challenge in RNA biochemistry. This toolkit enables future structural and cell-free studies through scalable access to native and engineered tRNAs, advancing the broader field of translation and synthetic biology.

## Introduction

Cell-free expression systems (CFES) are essential tools in biomanufacturing, medical and bioengineering research, and in foundational biology. The ability to synthesize proteins *in vitro* removes the constraints of living cells, enabling the expansion of the chemistries and functionalities of life. In biomanufacturing, these systems enable rapid and scalable production of proteins, enzymes, and therapeutics, including those that may be toxic or unstable in cellular environments^1^. For biomedical research, cell-free translation allows precise control over reaction conditions, enabling high-throughput screening, synthetic biology applications, and the development of novel diagnostics and vaccines^2^. In foundational science, it provides a simplified platform to study the fundamental mechanisms of gene expression, translation regulation, and protein folding, as well as enable the construction of synthetic cells^3^. In this context, tRNA serves as the essential adaptor substrate that links the nucleic acid sequence present in mRNA to the protein produced.

As mediators of protein expression, tRNAs dictate translational outcomes. Changes in the tRNA abundance can reprogram gene expression in response to stressors^4^, and can drive metastasis in cancer^5^. Viruses resupply degraded host tRNA pools during infection, enhancing the translation of their own proteomes^6^. Alterations of tRNA post-transcriptional modifications enact a diverse range of translational controls, such as initiating pathogenicity^7^, influencing embryogenesis^8^, or initiating mRNA decay through altered codon optimality^9^.

### tRNA modifications are essential to gene expression

tRNAs contain extensive post-transcriptional modifications critical to their function. The eponymous modifications of the D-loop and T-loop of tRNA (Dihydrouridine and riboThymidine, respectively) have been implicated in tRNA thermostability^10–12^. Pseudouridine synthase, TruB, aids in T-loop folding even if catalytically inactivated^13^. Nucleotides within the anticodon-stem loop (ASL) are extensively modified and bear the greatest chemical diversity^14^. These can range from either simple methylation or thiolation to so-called hypermodifications, which require several, often sequential enzymatic transformations^15^. ASL modifications are particularly important for tRNA function. In 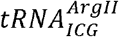, inosine-34 permits expanded wobble decoding towards CGU, CGC, and CGA codons^16^. In prokaryotes, I34 is formed by TadA, and is required in organisms which lack the tRNAs capable of decoding CGA codons^17–19^. Modifications at the 37-position of the ASL are also important for decoding efficiency and translocation. The hypermodified base 2-methylthio-6-pentenyl adenosine (ms2i6A) at A37 is necessary for efficient growth under stress^20^. Cyclic-threonylcarbamoyl adenosine (ct6A37) is an important positive determinant for the recognition of IleRS to 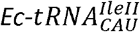^21^. In 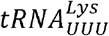, the combination of 5-methylaminomethyl-2-thiouridine (mnm5s2U) at U34 and the ct6A modification at A37 shapes the anticodon loop to permit efficient reading of the AAG codon in *E. coli*^*22*^.

Occasionally, ASL modifications can have dual roles. The methylated G37 (m1G) modification found among *Ec-tRNA*^*pro*^ isoacceptors is required for both efficient aminoacylation by ProS, and for maintaining translation open reading frames^23,24^. Similarly, the essential lysidine modification at C34 nucleotides enforces both correct aminoacylation and alternative wobble-decoding. Since 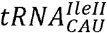 and 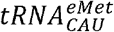 share a genomically encoded CAU anticodon, the lysidine-34 modification in 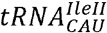 is required to maintain recognition by IleRS and enable wobble decoding of the AUA codon^25,26^.

### Generating tRNAs for *in vitro* translation

To generate tRNAs for *in vitro* studies, the standard technique is *in vitro* run-off transcription of linear tRNA templates using T7 polymerase^27^. While convenient and often high yielding, *in vitro* transcription using T7 polymerase requires a 5’-terminal guanine for efficient transcription initiation, and run-off transcription termination results in heterogenous 3’-ends^27^. To circumvent poor transcription initiation of tRNAs lacking 5’-guanine, 5’-leader sequences containing idealized T7 promoters are appended upstream of the tRNA which must be removed by subsequent cleavage by either self-cleaving tRNA-ribozyme fusions called transzymes^28–30^ or processing by RNase P^31,32^. In a similar fashion, 3’-ends can be made homogenous with either transzymes^33^ or the use of a 2’-OMe penultimate base^34^. Even after these challenges are addressed, tRNAs generated from *in vitro* transcription lack the post-transcriptional modifications critical to their function. Kinetic parameters determined using unmodified, *in vitro* tRNA substrates fail to recapitulate *in vivo* models of replication^35,36^, highlighting the limitations of such substrates in capturing physiological relevance.

After synthesis using *in vitro* transcription, post-transcriptional modifications can be added to tRNAs *in vitro*^*32*^, but the extensive range and chemical diversity of post-transcriptional modifications throughout tRNA requires numerous enzymes, substrates, and co-factors, making complete modification representative of *in vivo* tRNAs unrealistic. By contrast, tRNA sourced *in vivo* bear native modifications and properly processed 5’ and 3’ termini. Obtaining such tRNAs entails the isolation of the bulk tRNA pool from an organism of interest followed by the purification of individual tRNAs by either chromatographic separation of charged tRNAs using hydrophobic interaction chromatography or probe-based hybridization.

When separating tRNAs based on aminoacylation status, bulk tRNAs are first deacylated using a mild alkaline treatment and subsequently recharged *in vitro*.^*37,38*^. Charged tRNAs are then separated from the uncharged pool using hydrophobic interaction chromatography^38^. Such methods are often restricted to tRNAs charged with sufficiently hydrophobic amino acids to achieve reliable separation, although post-charging functionalization can expand this method to hydrophilic amino acids^39^. Still, challenges remain in separating individual tRNAs within isoacceptor families.

A more flexible method for tRNA purification relies on the sequence-specific hybridization of a target tRNA to a DNA oligonucleotide probe^40–43^. Here, an oligonucleotide probe is designed to specifically hybridize to a unique nucleotide sequence of the tRNA of interest while non-interacting tRNAs are washed away. Following the washes, the target tRNA is usually eluted by heat denaturation. By this technique, individual tRNAs can be readily separated from their sibling isoacceptors. Rarer tRNAs, in contrast, may struggle to outcompete their overly abundant, non-specific peers.

### tRNA overexpression enables purification of fully modified tRNAs

We sought to enable the production of fully modified tRNAs in high yield and in high purity. To do so, we coupled tRNA overexpression and isolation to DNA probe hybridization purification. We recovered target tRNAs in high yields (>10 nmol) with exceptional purity. Purified tRNAs retained functionality in cell-free translation reactions. We also show purification of non-native tRNAs. Specifically, we were able to express and purify 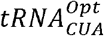 and 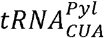 from the archaea *M. jannaschii* and *M. alvus*, respectively, and determined their modification status when expressed in *E. coli*. We found that modified 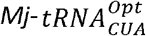 improved amber suppression in cell-free translation compared to its unmodified counterpart. We also demonstrate the purification of a mutant *E. coli* 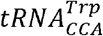 with a T-stem engineered to improve non-canonical amino acid incorporation.

This scalable and generalizable method produces *in vivo* modified tRNAs at high yield, enables access to both native and engineered tRNAs bearing complete post-transcriptional modification profiles, and overcomes a long-standing barrier in synthetic RNA biology. By streamlining tRNA preparation for structural, biochemical, and cell-free translation applications, this method stands to advance a broad range of efforts in protein translation and synthetic biology.

## Results and discussion

### Purification of native E. coli tRNAs using selective overexpression

Previous work has suggested that oligonucleotide probe purification would benefit from greater abundance of target tRNA, improving both yield and purity^43^. To this end, we cloned and overexpressed 10 native *E. coli* tRNAs using a previously described expression vector^44^. Overexpressed tRNAs were isolated using acid-phenol extraction and sequential isopropanol precipitation, yielding enriched tRNA pools ranging from 5-10 mg from 1 L of culture (**Figure 1a**). tRNAs of interest were clearly overexpressed **(Figure 1b)**, and target tRNAs were successfully purified using columns with oligonucleotide probes designed to hybridize specifically to each tRNA (**Materials and Methods**).

**Figure 1.**
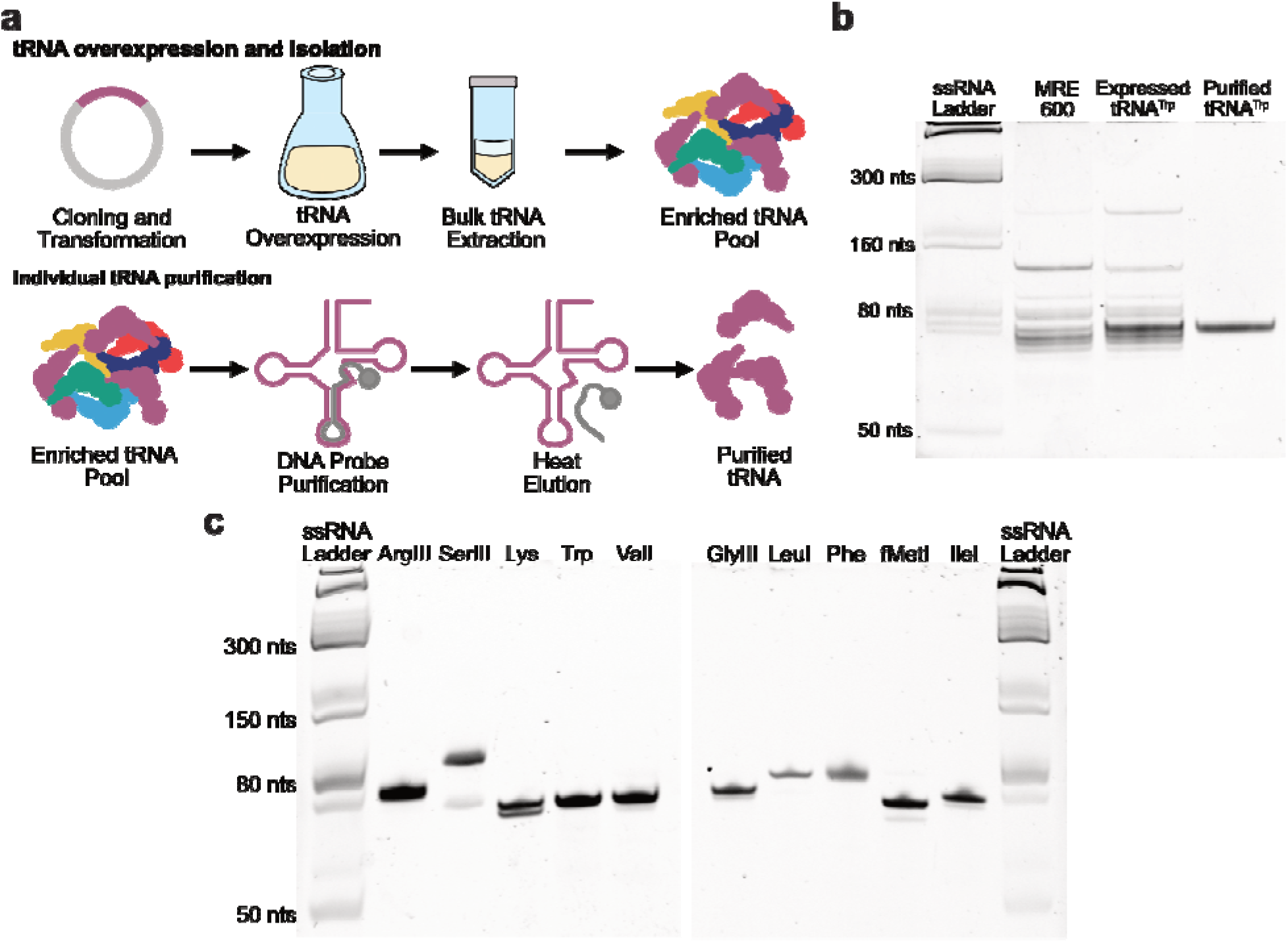
Purification of Native *E. coli* tRNAs. **(a)** Method of overexpression and purification of individual *E. coli* tRNAs. Target tRNA is overexpressed in *E. coli* and the bulk tRNA pool enriched with the target tRNA is recovered using acid-phenol extraction. After extraction, enriched target tRNA is bound to a column coupled to a DNA hybridizing probe specific to the overexpressed tRNA. Non-specific tRNAs are washed away, and target tRNAs are heat eluted to produce highly pure tRNA. **(b)** Urea-PAGE of overexpressed and *in vivo Ec*-purified using DNA-probe hybridization column purification. Commercial MRE600 serves as an expression control. **(c)** Urea-PAGE of each native *in vivo E. coli* tRNA purified in this study.

Purity of recovered tRNAs was first determined using denaturing urea-polyacrylamide gel electrophoresis (urea-PAGE). Single bands were observed for five of the ten purified tRNAs while faint, smaller molecular weight bands were observed for the remaining tRNAs (**Figure 1c**). These fainter bands are likely degradation products or partially folded tRNA structures that persist despite running the gels under denaturing conditions.

To further confirm the identity of the recovered tRNAs, we sought to characterize them by analyzing their signature digestion products (SDPs) resulting from treatment by RNase T1 digestion using MALDI-TOF^45^. Specifically, we were interested in confirming SDPs for target tRNAs, as well as determining the presence of any potential contaminating tRNA. Indeed, all purified tRNAs contained neither unique fragments of their sibling tRNA isoacceptors, nor any SDPs from other *E. coli* tRNAs (**Supplemental Data File 1**).

### Intact LC-MS confirms modification and purity

To further confirm the purity of tRNAs, as well as gain deeper insights into their modification status, we utilized intact tRNA LC-MS, a recently developed technique employed to quantify tRNA aminoacylation^46^. As aminoacylation is a form of transient post-transcriptional modification, we hypothesized intact tRNA LC-MS could resolve different modification states of purified tRNAs.

Each tRNA eluted as a single major peak or a major and minor peak (**Figure 2**). For each tRNA, masses were assigned according to literature reported modification profiles found in the MODOMICS database^47^ (https://genesilico.pl/modomics/). An exception to literature reported masses were tRNAs reported to contain threonylcarbamoyl adenosine (t6A37). *In vivo*, fully modified *E. coli* tRNAs normally contain ct6A37, while the MODOMICs reported t6A37 is an artifact of earlier efforts of modification profiling^48,49^. Accordingly, calculated masses for the 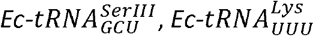, and 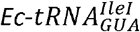 were assigned using ct6A37.

**Figure 2.**
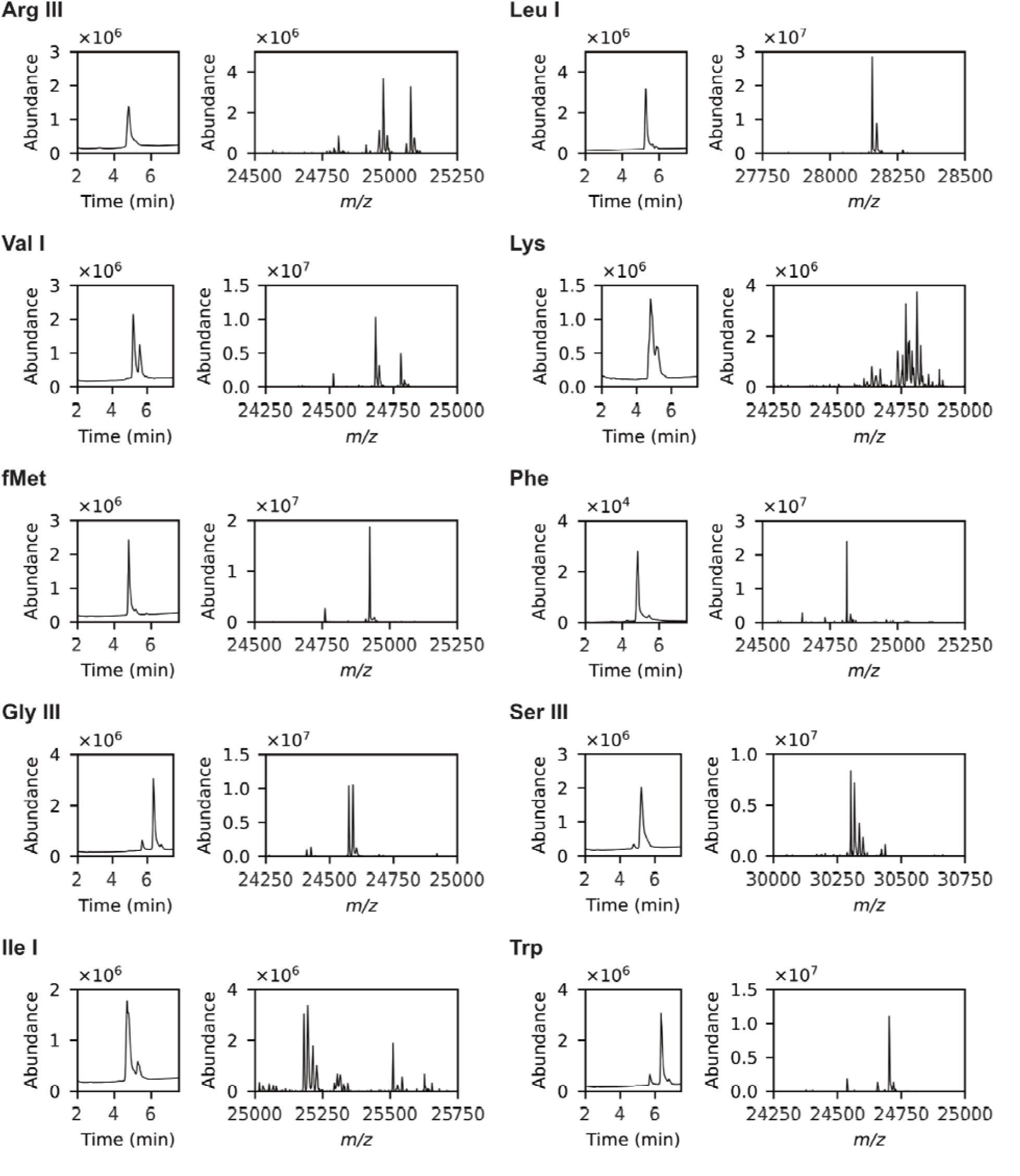
Intact tRNA LC-MS of purified native *E. coli* tRNAs. Ten purified native tRNAs from *E. coli* were analyzed using intact tRNA LC-MS. For each tRNA, left plots represent total ion chromatograms of each tRNA and right plots represent deconvoluted, extracted ion chromatograms of individual tRNAs. Most tRNAs eluted as a single peak. Extracted ion chromatograms reveal individual modification states of each purified tRNA.

Intact tRNA LC-MS revealed a range of modification states not reported in MODOMICS. Across all tRNAs species, we observed masses either 14, 16, or 30 *m/z* greater than those found in MODOMICS (**Supplemental Data File 2**). Considering these are mass shifts corresponding to additional methylation (14 m/z), thiolation (16 m/z), or both, we suspected modification enzymes may be recruited to overexpressed tRNA normally outside their substrate scope.

In prokaryotes, the invariant U8 is thiolated by ThiI across nearly all tRNAs, generating 4sU8^50,51^. U8 thiolation is dynamic and changes throughout growth^52^. Recently, a method used to detect 4sU8 using RNA-seq uncovered 4sU8 modifications within tRNAs that were not reported in either MODOMICS or tRNAdb^53^. We find these thiolation modifications as well and identify 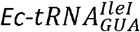 as another candidate for 4sU8 modification.

Broad methylation is best explained by the tRNA methyltransferase, TrmH, which modifies G18 among 14 of 46 tRNAs in *E. coli*^54^. Like thiolation at U8, G18 methylation (Gm18) is dynamic and changes due to organismal stress^55^. Of the native tRNAs purified from this study, only 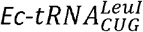 is expected to contain the Gm18 modification^47^. We surmise that overexpression has made the other purified tRNAs competent for methylation. For example, it was recently shown that 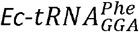 is partially methylated by TrmH despite not being considered a canonical substrate^54^.

Intact tRNA LC-MS revealed unknown modifications for the rare tRNA 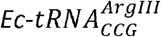^56^. Like the other tRNAs studied, 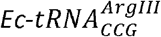 contained species consistent with additional thiolation and methylation when compared to its MODOMICS reported mass. Unexpectedly, we observed three peaks ~100 m/z above the reported mass of 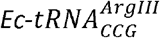 and the observed thiolated/methylated variants (**Figure 3a**). This mass shift is likely due to modification of uridine-47 to 3-amino-3-carboxypropyl-uridine 47 (acp3U47), a unique modification shared among a small group of tRNAs. These tRNAs include 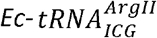, the sibling isoacceptor to 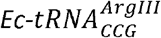 While 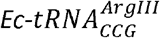 contains the requisite 7-methylguanosine modification (m7G) at G46 required for acp3U47 modification^57^, MODOMICS reports the unmodified U47 instead^47^. Recently, Wang et al. exploited the tendency for modifications to abort reverse transcription to profile modifications using deep-sequencing across *E. coli* tRNAs^58^. They found acp3U47 reliably terminated transcription and confirmed the acp3U modification in 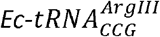 ^58^. We share these findings and conclude that the MODOMICS reported modification profiles for rarer tRNAs like 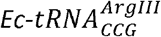 might be incomplete.

**Figure 3.**
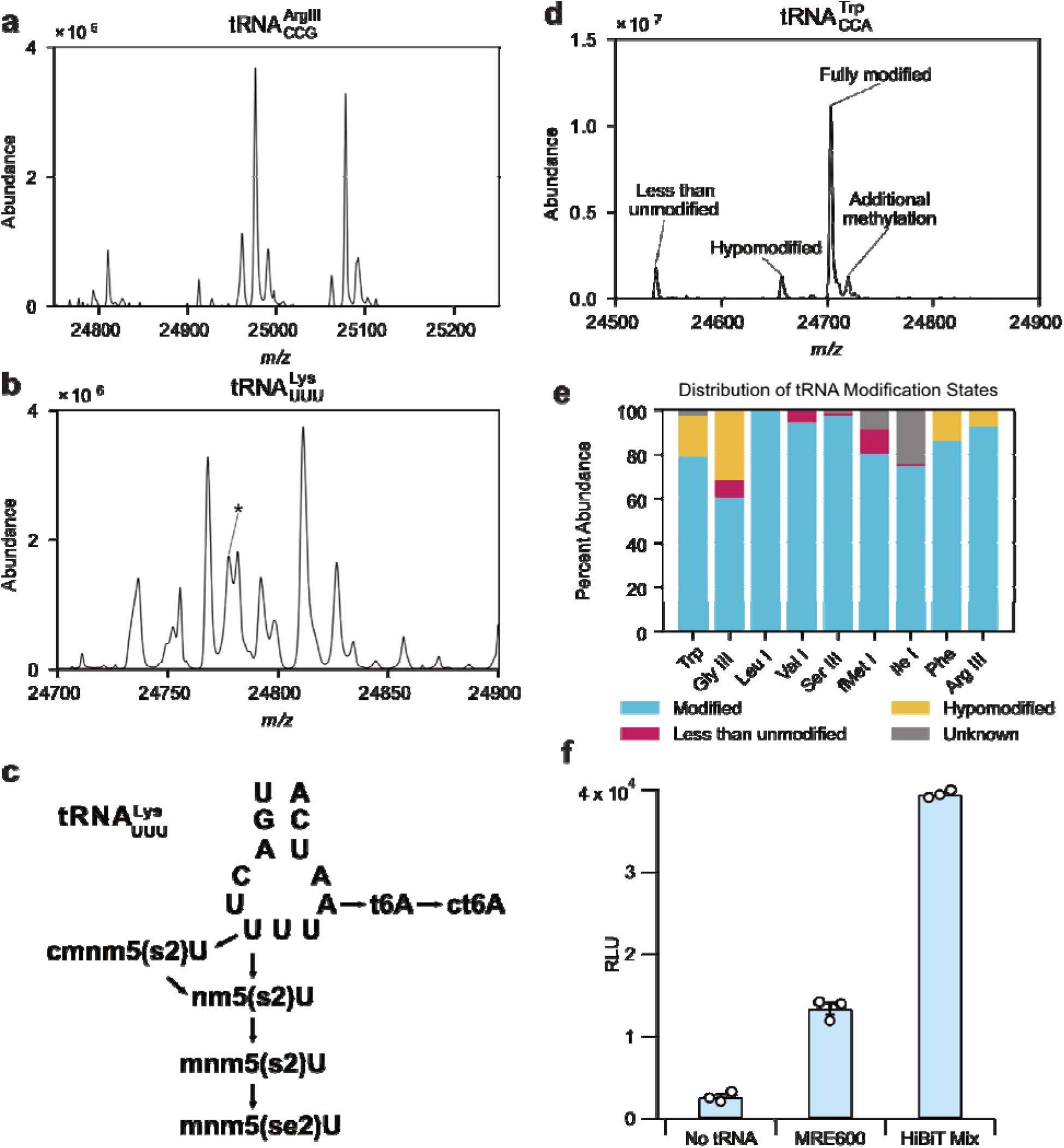
Intact tRNA LC-MS reveals diverse modification states of tRNA. **(a)** *Ec*-LC-MS extracted ion chromatograms containing a doublet of triplet peaks where the triplets are separated by ~100 m/z. **(b)** Extracted ion chromatogram of *Ec*-shows a plurality of modifications. Peak corresponding to the MODOMICS reported *Ec*-modification profile is labeled with an asterisk (*). **(c)** Modification intermediates within the tRNA^Lys^ anticodon stem loop demonstrate the plurality of potential 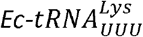 isoforms. U34 undergoes a series of modifications. First, U34 is modified to either nm5U or cmnm5U by the MnmE:MnmG complex depending on the substrate used. Then, cmnm5U is converted to nm5U by MnmC. Whether formed by MnmE:MnmG or MnmC, nmn5U is methylated by MnmC to generate mnm5U. At any stage of modification, the 2-carbonyl oxygen of uridine can be thiolated to 2-thiouridine. If available, mnm5s2U34 can have its sulfur exchanged to selenium. Beyond U34 modifications, A37 is modified to ct6A from an t6A intermediate which is susceptible to nucleophilic attack. **(d)** Representative categorization scheme of extracted ion chromatogram of 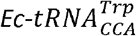 “Less than unmodified” categorization represents an ion species with a mass less than if the target tRNA was completely unmodified, suggesting a potential tRNA degradation product. “Hypomodified” ion species correspond to tRNA with an identified intermediate modification; here, hypomodified 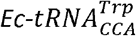 contains i6A37 rather than the fully modified ms2i6A modification. “Modified”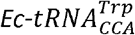 includes both MODOMICS reported mass along with an additionally methylated isoform identified from intact LC-MS. **(e)** Percent abundances of each tRNA species identified using intact LC-MS as categorized in **d. (f)** Steady-state luminescence maxima of NanoGlo reactions completed by HiBiT peptide translated using tRNA pools. Bars represent the average of steady-state, luminescence maxima. Open circles represent experimental replicates. Error bars represent error as standard deviation.

The most difficult to assign species of tRNAs were those containing ct6A37 within the anticodon loop: 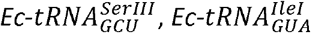, and 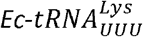. Along with putative methylations, 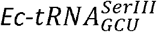 and 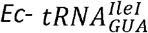 contained masses ~32 m/z greater than expected. These are most easily described by an alternative modification pathway of t6A37. Rather than being cyclized TcdA, t6A37 is methylated by the tRNA methyltransferase, TrmO, generating 6-methythreonyladenosine (m6t6A). In *E. coli* m6t6A37 modification is exclusive to 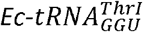 and 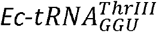, where TrmO relies on the C31:G39 pair and G34 as major positive identity elements for recognition^59^. 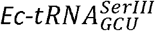 and 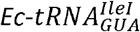 both contain these elements, but the absence of G35 base should preclude their modification by TrmO^60^. Still, overexpression may be artifactually recruiting TrmO. Another possibility is the formation of an amide adduct. The hydantoin ring of ct6A37 is susceptible to nucleophilic attack by primary amines^48,49^, such as methylamine, which could describe this mass shift.

A diverse collection of masses was observed for 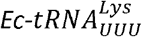 making even partial mass assignments impractical. We were only able to confidently assign the MODOMICs reported mass of 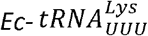 (**Figure 3b**). Surprisingly, this was not the most abundant ion species detected; instead, an ion product 34 m/z greater had the greatest abundance. The inability to assign complete masses for 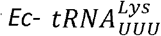 may stem from the complex and dynamic nature of its unique ASL modifications **(Figure 3c)**. Modification of 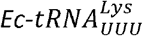 is first initiated by MnmE and MnmG which, in concert, form either 5-carboxylmethylamino (cmn5U) or 5-methylamino uridine (mn5U) ^60^. MnmC subsequently catalyzes the cleavage of the carboxymethyl group, if present, and stimulates the subsequent methylation of the aminomethyl group, forming 5-methylaminomethyl uridine (mnm5U). These modification states can vary depending on the growth conditions^61^, and it is unclear whether overexpression may bias towards one modification state. In addition to modifications at the 5C of uridine-34 of 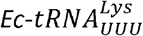, the 2C carbonyl oxygen is thiolated independently by MnmA to form 5-methylaminomethyl-2-thiouridine (mnm5s2U). If selenium is present, the 2-thiol group within mnm5s2U can be exchanged for a selenium atom through a geranyl intermediate, thereby rendering the end-stage hypermodification product, mnm5se2U^62,63^. Thus, the multitude of modifications at U34 along with the controvertible modification state of ct6A37 can generate an array of potential modifications for 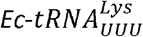 that we were unable to reconcile.

We determined the relative abundances of each tRNA modification isoform within a sample according to peak area. To assign relative abundances of tRNA modification states, we categorized tRNA masses as either less than unmodified (i.e., masses that are lesser than if the tRNA lacked any modification at all, perhaps as the result of degradation), hypomodified, fully modified/alternatively modified (i.e. containing masses consistent with literature reported masses and/or additional methylation, thiolation, etc.), or unknown (i.e., masses greater than fully modified or acylated tRNA, likely pre-mature tRNA still containing leader or trailer sequences). An example of assignment is shown in Figure 3c for 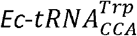.

Purified tRNAs contained mostly fully modified or alternatively modified species (**Figure 3e**). Overexpression seemed to subject tRNAs to a greater level of modification, gaining modifications that are not reported in MODOMICS. Still, there were a few instances of hypomodification. For example, 9% of 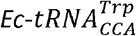 isoforms contained i6A37, the intermediate modification to ms2i6A37 (**Figure 3e; Supplemental Data File 2**). Incomplete modification here is problematic as tRNAs lacking the 2-methylthio moiety exhibit 3-9 fold increase in ribosomal frameshifting^64^.

Additionally, roughly 30% of the 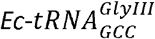 species were found to be lacking a thiolation modification, likely 4sU8. Unlike the incomplete modification of ms2i6A, the absence of this modification is expected to be less detrimental. For certain tRNAs, absence of U8 thiolation results in poorer thermostability and can result in degradation *in vivo*, though 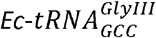 was not shown to be impacted^65^. The extent of 4sU8 modification varies in 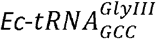 even at native expression levels^53^. Aside from these isolated cases of hypomodification, these results show that our method yields highly pure tRNA, with the vast majority carrying their complete modification profiles.

### Recovered tRNAs are competent for translation

Following successful purification, we wanted to confirm that purified tRNAs could still participate in translation. To test this, we pooled together 10 unique tRNAs required to translate HiBiT, an 11-amino acid reporter peptide, that binds with LgBit to form a functional NanoLuc enzyme^66^. Each purified tRNA was supplied in equimolar amounts, such that each comprised one-tenth of the total tRNA pool. This designed pool then served as the tRNA complement to a ΔtRNA PURE cell-free expression reaction tasked with the translation of the HiBiT reporter peptide. Strikingly, the purified tRNA pool rescued HiBiT translation (**Figure 3f**) and outperformed the complete MRE600 tRNA pool typically used to complete PURE^67^ by roughly 4-fold.

### Purification and Modification Profiling of 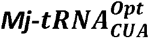

A key goal in synthetic biology is genetic code expansion, which adds new chemistries to the codon table. tRNAs are critical to these efforts, but the modification state of the tRNAs used in genetic code expansion are largely unexplored. We therefore sought to expand the scope of our method to archaeal tRNAs used for genetic code expansion expressed in *E*.*coli*. We began by investigating 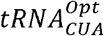 from *Methanocaldococcus jannaschii*, the tRNA partner to several *Mj*TyrRS mutants capable of introducing a wide array of tyrosine analogs into translated proteins^69^ and whose modification patterns have already been partially characterized^69^.

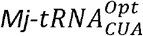 expression led to substantial tRNA enrichment (**Supplemental Figure 1**), and we proceeded to purify 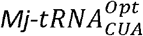 using a probe designed to bind to its anticodon. After several bindings, the amount of 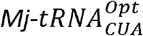 recovered from the column did not change (3-4 nmoles per binding), showing purification yields are limited by the binding capacity of the column. To improve upon this, we prepared another column, increasing the amount of probe coupled to the resin 5-fold. The resulting column increased purification yield to 10 nmoles, exhausting the 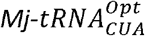 aliquot after two consecutive bindings. The purified 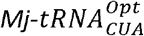 ran as a single band on urea-PAGE (**Supplemental Figure 1**), consistent with high purity.

It was previously reported that 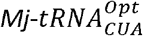 is modified by the native modification machinery of *E. coli*^*69*^. We sought to confirm these modifications and attempt to complete its modification profile. First, we used Signature Digestion Product (SDP) analysis using MALDI-TOF. To generate an initial fragment profile of 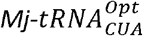, we performed RNAse T1 digestion of 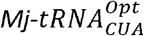 *in silico* devoid of any modifications using the Mongo Oligo Mass Calculator (http://rna.rega.kuleuven.be/masspec/mongo.html). We were able to assign 9/12 unmodified digestion fragments detectable to our MALDI-TOF method using this initial catalog of masses (**Supplemental Figure 2a-b**). The remaining three fragments generated from *in silico* digestion encompass regions that are commonly modified in *E. coli*: U8, the anticodon loop, and the TΨC loop. Thus, we generated a new draft sequence of 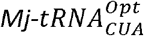 for *in silico* digestion where: 1) U8 is modified to 4-thiouridine, 2) U54 is methylated to ribothymidine, and the inclusion of the ms2i6A modification at A37, as first reported by Baldridge et al^69^. Doing so allowed us to identify masses for 4-thiouridine (4AGp), rT54 (TUCAAAUCCCCUCCGp), and ms2i6A37 (ACUCUA*AUCCGp) within the MALDI spectrum.

We were surprised to see a set of 4 masses clustered around where ACUCUA*AUCCGp was expected (**Supplemental Figure 2b**). Since ms2i6A is a hypermodified base within this fragment, we considered if any of the masses could be described by hypomodification. We were able to identify the masses which correspond to the intermediate i6A modification. Finally, the other two masses are best explained by a methylation of ACUCUAAAUCCGp fragments containing either ms2i6A or i6A modifications. The anticodon loop of 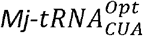 closely matches *E. coli* tRNA^Trp^, which along with ms2i6A37, contains a methylated 2’-OH of C32, suggesting that 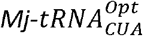 might also contain a methylated C32.

To decode the full sequence and modification profile of 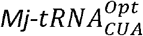, we performed Next-Generation Mass Spectrometry sequencing (NGMS-Seq)^70–72^ (**Figure 4**). Corroborating our MALDI-TOF analysis, five modifications were identified at three positions, including partial modifications at position 32 (C/Cm: 82%/18%), 37 (i6A/ms2i6A: 84%/16%), and 46 (G/mG: 91%/9%) (**Figure 4a-b**). These results are supported at both the intact RNA level (pre-hydrolysis) and fragment level (post-hydrolysis). Homology searches of intact masses revealed five isoforms differing by as little as a single nucleotide or modification (**Figure 4c**) and potential partial modifications. For example, a 14 Da difference between IF1 and IF5 indicates partial methylation, later confirmed at position 46 (G/mG) by a distinct 2D mass–retention time (t_R_) ladder branch.

**Figure 4.**
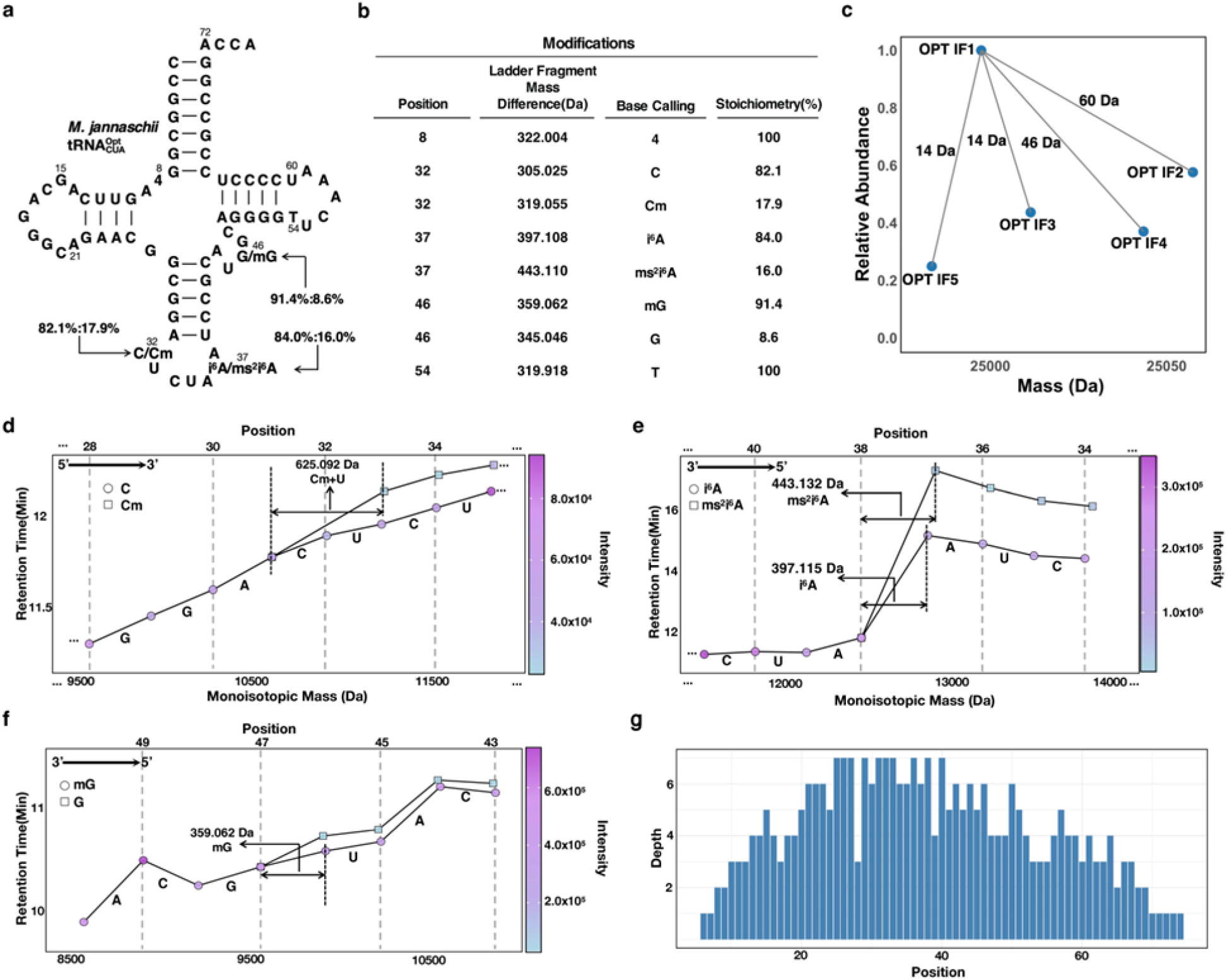
NGMS sequencing decodes the complete sequence and modification profile of *Mj*-, including partial modifications and stoichiometry at position 32, 37, and 46. **(a)** De novo base calling of the full-length tRNA sequence, identifying three partial tRNA nucleotide modification positions with distinct stoichiometries: C/Cm (82%/18%) at position 32, i^6^A/ms^2^i^6^A (84%/16%) at position 37, and G/mG (91%/9%) at position 46. **(b)** Positions and stoichiometries of noncanonical and partially modified nucleotides in *Mj*-. **(c)** Homology search of intact tRNA masses revealed coexisting isoforms that may differ as little as one nucleotide or modification. For example, a 14 Da difference between IF1 and IF5 indicates partial methylation, which was further confirmed by NGMS-Seq to be G/mG at position 46. **(d)** a 2D mass-t_R_ branch from the original 511-ladder at position of 32 indicates a partial methylation (Cm) at this position, confirms coexisting Cm and C (82.1%/17.9%). A 511-ladder fragment is missing at position 32 because the 2⍰-O methyl group of Cm blocks formic acid hydrolysis of the phosphodiester bond between itself and the following nucleotide U, thus creating a mass gap of 625 Da, which corresponds to Cm+U. **(e)** A 2D mass–t_R_ branch from the original 311-ladder at position 37 confirms coexisting i^6^A and ms^2^i^6^A (84.0%/16.0%). **(f)** A 2D mass–t branch from the original 3⍰-ladder at position 46 confirms partial methylation (G/mG: 91.4%/8.6%). (g) Sequence depth-position plot shows high read coverage across positions, supporting the reliability of NGMS-Seq through multiple ladder confirmations. Depth indicates the number of times each nucleotide position was read independently from different tRNA ladder or adduct forms.

In 2D mass–t_R_ plots, each partial modification produced a distinct branch diverging from the original 3⍰ or 5⍰ ladder that carries the canonical nucleotide counterpart, enabling precise identification, localization, and stoichiometry determination of the partial nucleotide modification. Analysis of hydrolyzed fragments showed distinct 2D mass–t_R_ branches at positions 32, 37, and 46, each corresponding to a specific partial modification (**Figure 4d-f**). The stoichiometry at each site was determined using relative ladder fragment intensities of modified versus unmodified ladder fragments. For instance, the stoichiometry at position 46 (G/mG) was calculated as 91%/9% based on relative MS intensity ratios of ladder fragments containing canonical G and modified mG at the position. At position 32, the 2⍰-O-methyl group of Cm blocked formic acid hydrolysis between Cm and the following U, producing a 625 Da mass gap (Cm+U) and confirming the modification. The final stoichiometry was calculated as 82.1% C and 17.9% Cm. A sequence depth-position plot (**Figure 4f**) showed consistently high read coverage across all positions, further validating NGMS-Seq’s reliability.

Intact LC-MS and assembled masses from NGMS-seq gives an insight into the stages of modification and their abundances in the expression of 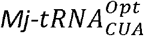. The isoform with the greatest mass and second greatest abundance (OPT IF2, **Figure 4c**) included the modifications Cm32 and ms2i6A37. No further modifications were identified beyond this isoform, and it can be considered the mature isoform of the tRNA. In contrast, the most abundant isoform (OPT IF1, **Figure 4c**) contained only i6A37 within the anticodon loop, suggesting subsequent methylation of C32 and methylthiolation of i6A37 may be rate limiting steps in 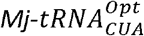 modification. No isoform identified contained an unmodified A37 suggesting that prenylation by MiaA to form i6A occurs rapidly.

### Modified 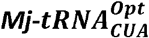 decodes UAG stop codons more efficiently than unmodified 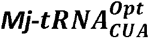

We wanted to assess how *in vivo* purified 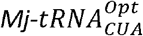 performed in amber suppression assays compared to unmodified 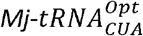. To do so, we translated two different amber suppression reporters: a HiBiT:LgBit split luciferase system encoding an an amber codon at the third position upstream of HiBiT peptide and a previously established super-folding GFP where the T216 residue is mutated to an amber stop codon (sfGFP216TAG)^73^. These templates were used in a PURE cell-free translation reactions containing either *in vivo* purified or *in vitro* transcribed 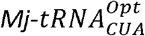 along with their partner aminoacyl tRNA synthase (AARS), pCNFRS^74^, and *para*-cyanophenylalanine as the aminoacyl substrate.

When suppressing sfGFP216TAG, unmodified 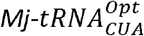 failed to translate sfGFP sufficiently to be distinguishable from background (**Figure 5a**). Similarly, for HiBiT translation, modified 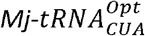 produced 2-fold greater HiBiT compared to unmodified 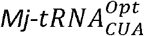. To account for potential misincorporation resulting in false positives from these read-through assays, we subjected the HiBiT translation reactions to FLAG-tag purification and analyzed the purified peptides using MALDI-TOF. Both modified and unmodified 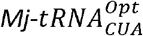 produced peaks consistent with incorporation of para-cyanophenylalanine (**Figure 5b**). There was no significant misincorporation observed from peptides produced using either *in vivo* or *in vitro* 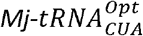 (**Figure 5b**). These results demonstrate that the post-transcriptional modifications of *in vivo* purified 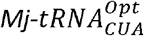 are critical for amber suppression.

**Figure 5.**
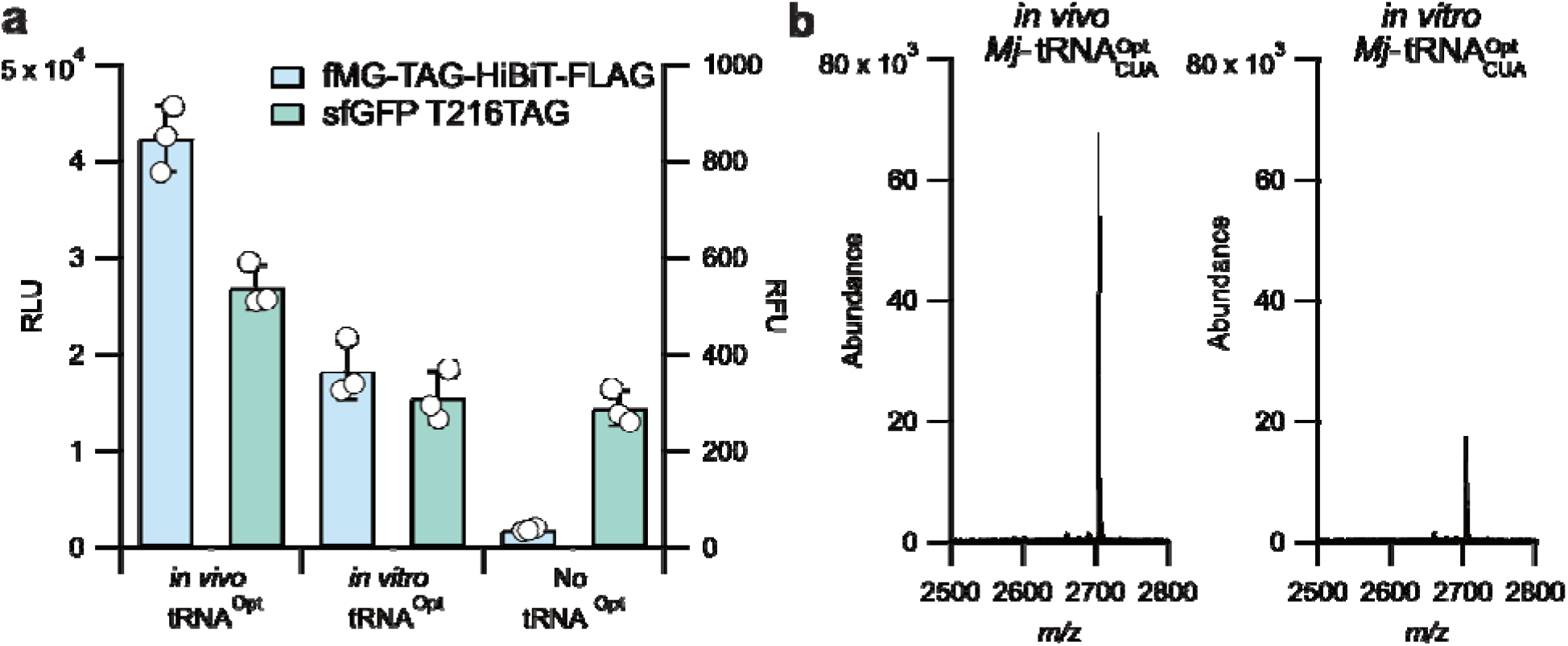
Cell-free amber suppression is improved using *in vivo* purified *Mj*-. **(a)** Yields of cell-free expression of HiBiT or sfGFP amber suppression reporters mediated by either *in vivo* or in vitro generated *Mj*-. No added *Mj*-served as a negative control. Blue bars (left y-axis) represent the average of steady-state, luminescence maxima from NanoGlo reactions following PURE cell-free expression. Green bars (right y-axis) represent the average end-point fluorescence of sfGFP translation. Open circles represent experimental replicates. Error bars represent error as standard deviation. **(b)** MALDI-TOF of amber suppressed HiBiT peptides purified from PURE CFES amber suppression reaction from **a)**. Calculated mass of [fMG-pCNF-HiBiT + H]^+^ = 2703.287 *m/z* and observed masses of 2703.434 and 2703.402 from *in vivo Mj*- and *in vitro Mj*-suppressed reactions, respectively.

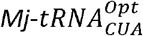 is the product of extensive engineering of its parent tRNA, *Mj-*tRNA^Tyr^. In addition to mutations in its acceptor stem and T-stem for increased affinity to EF-Tu^75^, mutational screens across the anticodon loop yielded a G37A mutation^76^ resulting in an anticodon loop competent for MiaA modification^77^. Similar mutational studies have rendered anticodon loops capable of ms2i6A modification^78,79^. While not essential, ms2i6A has long been established as important for efficient translation. The loss of either ms2i6A or i6A modifications generate tRNAs with reduced affinity towards ribosomes^80^ and prevents the production of virulence factors in *Shigella flexneri*^*81*^. Similarly, *E. coli* lacking *miaA* were incapable of decoding UAG codons^69^, indicating UAG decoding in *E. coli* is linked to ms2i6A modification.

It is uncertain if these prokaryotic attunements faithfully mimic the archaeal anticodon loop structure or are necessary to coerce non-native tRNAs to function in unrelated hosts. The i6A37 and ms2i6A37 modifications are absent in Archaea, which instead use either threonylcarbamoyl or hydroxynorvalylcarbamoyl modifications at A37 if they are modified at all^82–84^. Moreover, the native progenitor tRNA of 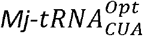 contains a G37 which is normally hypermodified to demethylwyosine (imG14) in *M. jannaschii*^*85*^. Thus, the inclusion of ms2i6A37 or i6A37 likely represents a significant departure from the native archaeal anticodon loop topology. As a greater set of archaeal tRNA and AARS pairs are considered for codon table expansion^86,87^, it may be important to address the dual challenge of aligning heterologous host requirements to proper tRNA anticodon loop structure.

### Modification Profiling of 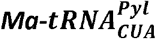

We were additionally interested in purifying 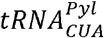 from *Candidatus methanomethylophilus alvus*, the tRNA partner of *Ma*PylRS. This tRNA/AARS pair has garnered great interest due to its exceptional substrate scope, including diverse side-chains and non-canonical backbones^88,89^. Surprisingly, 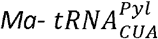 failed to overexpress as robustly as native *E. coli* tRNAs or 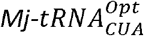 despite using the same pKK223-3 backbone (**Supplemental Figure 3a**). Chemiluminescent northern blotting revealed that 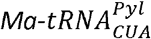 was indeed expressed but poorly (**Supplemental Figure 3b**). When the expressed tRNA isolate was used in purification, several native *E. coli* tRNAs were co-purified along with target 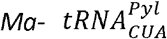 (**Supplemental Figure 3c**). This observation emphasizes the importance of tRNA overexpression for the successful purification of tRNAs in high yield. Since the pKK223-3 backbone maintains a low copy number, we considered if switching to a higher copy number plasmid would improve the expression yield of 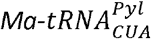. We also considered if leaky expression from the lac repressor system might contribute to reduced expression yields due to toxic background expression. While it is unclear if uncharged tRNAs are subjected to degradation^90,91^, we also hypothesized that co-expression of *Ma*PylRS with 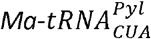 in the presence of the cognate amino acid substrate, Boc-Lys, would further improve purification yields.

Correspondingly, we designed a new, fully synthetic plasmid expression construct that incorporated the high copy number pUC19 origin of replication where 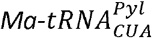 is under the control of a Pm:XylS positive transcriptional activating system^92^. *Ma*PylRS was also included within the vector under arabinose induction. The resulting constructs overexpressed the target tRNA and enabled purification of 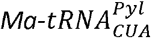 (**Supplemental Figure 4**). Interestingly, along with a band at the expected 71 nts for 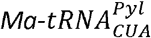, we also observed a larger band at roughly 80 nts, potentially indicative of incomplete processing (**Supplemental Figure 4**).

We were interested in cataloging the modifications of 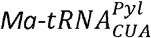. We were especially interested in the modification state at A37. We used MALDI-TOF to generate a draft profile of 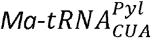 modifications and attempt to identify the co-purifying tRNA. We were able to assign a singular SDP relating to endogenous native 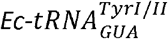, but were unable to match any other 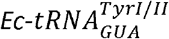 SDPs indicating the 80 nt band observed using urea-PAGE might in fact be an immature 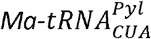 processing intermediate. We were able to assign 7/9 fragments that would result from the RNAse T1 digestion of unmodified 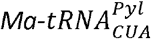 **(Supplemental Figure 5a-b)**. The remaining unassigned, unmodified fragments were again found to be within the typically modified anticodon and T-loops, suggesting the presence of post-transcriptional modifications. We were able to identify fragments when including modifications for ms2i6A37 and rT54 **(Supplemental Figure 5a-b)**. Interestingly, we failed to observe any evidence of the C32 methylation found in 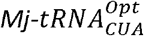.

We proceeded with NGMS-seq to obtain stoichiometric and sequence-specific modification data. To do so, we first performed an additional purification step using hydrophobic interaction chromatography to separate purified 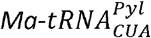 from higher molecular weight contaminants. HIC separated the purified 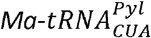 into three fractions (**Supplemental Figure 6a**). Urea-PAGE revealed Fraction 3 contained the correct length 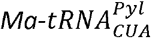 (**Supplemental Figure 6b**) which was used for characterization using NGMS-seq. NGMS-Seq of 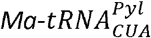 revealed full sequence and partial modification stoichiometry at position 37, identifying three coexisting forms and their stoichiometries: A (23.4%), i6A (13.8%), and ms2i6A (62.8%) (**Figure 5**). Intact mass differences (46 Da and 114 Da) supported these findings prior to hydrolysis and were validated by 2D mass–t_R_ ladder analysis. Branching at position 37 confirmed the partial modifications, with stoichiometry derived from fragment intensities. High sequence depth further validated the accuracy and reproducibility of NGMS-Seq.

**Figure 6.**
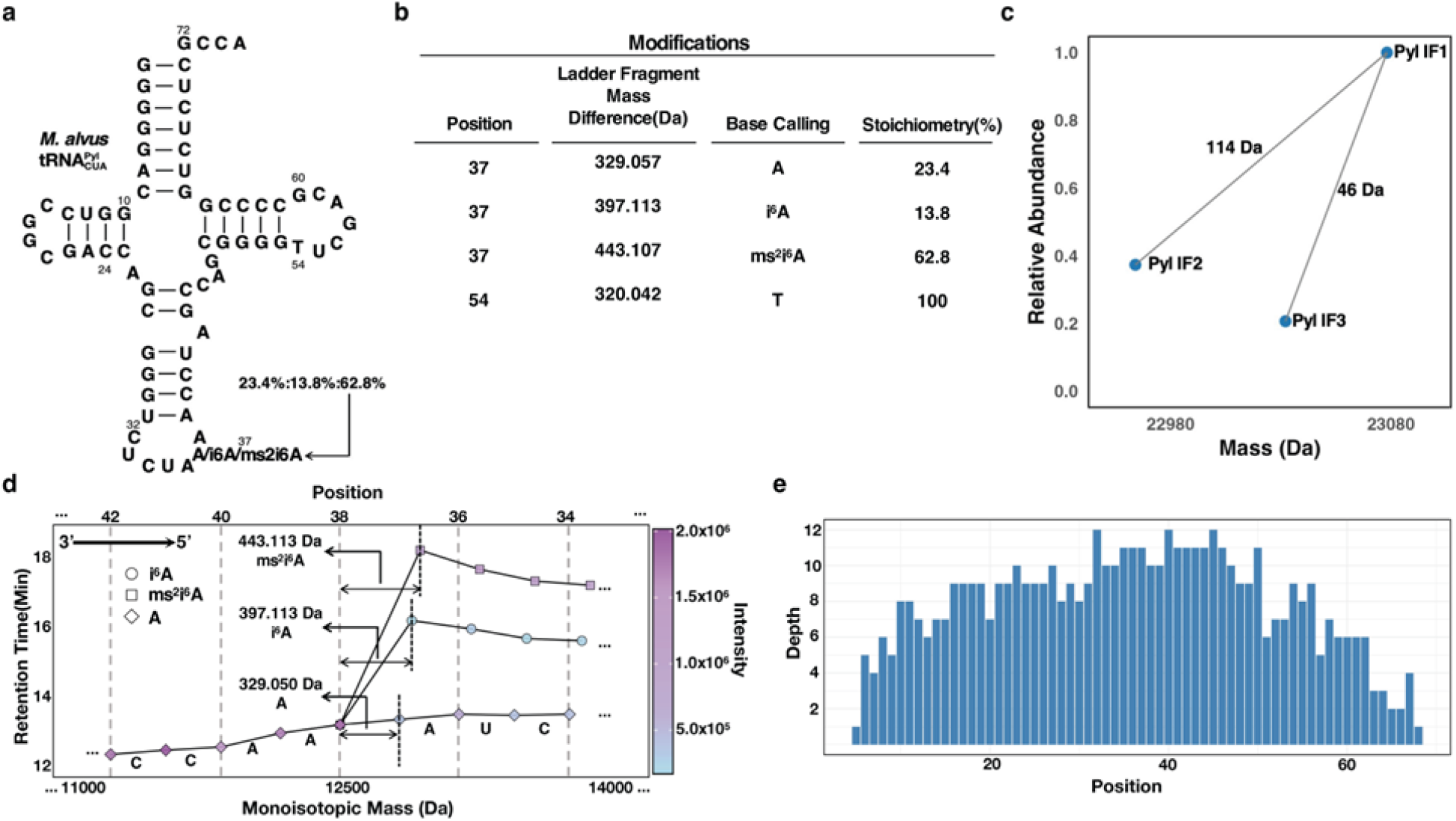
NGMS sequencing decodes the complete sequence and modification profile of *Ma*-, including partial modifications and stoichiometry at position 37. **(a)** De novo base calling of the full-length tRNA sequence, identifying partial tRNA nucleotide modifications at position 37 with stoichiometry: 23.4%/13.8%/62.8% (A/i^6^A/ms^2^i^6^A). **(b)** Position and stoichiometry of the noncanonical nucleotides and partially modified nucleotides in ***Ma***-. **(c)** Homology search of intact tRNA masses identifies coexisting isoforms differing by single modifications. A 46 Da difference between IF1 and IF3 suggests partial modification between i6A and ms2i6A; a 114 Da difference between IF1 and IF2 indicates coexistence of A and ms2i6A. NGMS-Seq confirms these differences at position 37. **(d)** 2D mass–tR branches at position 37 confirm the presence of A, i6A, and ms2i6A. Stoichiometry was determined using fragment intensity ratios, as described in Method Section. **(e**) Sequence depth-position plot shows strong coverage across all positions, supporting NGMS-Seq reliability through multiple independent ladder reads.

Unlike wild-type 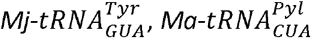 contains the MiaA recognition sequence natively, though as noted previously, archaea lack *miaA* and *miaB* needed to confer the i6A and ms2i6A modifications. While the native modifications of 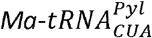 are not known, the anticodon loop of another archaeal methanophile, *Methanosarcina barkerii* 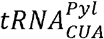, is unmodified. For *M. alvus*, the inherent flexibility of the tRNA^Pyl^ body seems to be critical for its activity^93^ and might be aided by additional flexibility conferred by ms2i6A within the anticodon loop^94^. Like 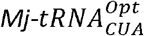 subsequent studies will have to scrutinize the role ms2i6A plays in translation efficiency for 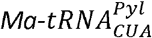 or if alternative modifications might further improve its decoding activity.

### Purification of an Engineered 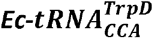

In a final demonstration of our system, we wanted to purify mutant *E. coli* tRNAs that may improve the ribosomal incorporation of ncAAs. A common strategy in ncAA incorporation is to exploit the substrate promiscuity of native *E*. coli AARSs^95^. Once charged, these tRNAs are delivered by EF-Tu to the elongating ribosome. EF-Tu binds charged tRNAs at both the acceptor and T-stems such that there is a thermodynamic balance in affinity^96^. As such, tRNA T-stems are matched to their aminoacyl cargos to achieve a uniform affinity^97,98^. Promiscuous acylation can challenge thermodynamic matching by coupling challenging aminoacyl substrates to weakly interacting T-stems. One solution to this challenge is to utilize tRNAs which contain high affinity T-stems that compensate for the poor binding at the amino acid binding pocket^99–101^.

To this end, we mutated the native *E. coli* 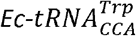 to contain the T-stem of *Ec-*tRNA^*Asp*^. Both *Ec-* tRNA^*Asp*^and *Ec-*tRNA^*Glu*^require a high affinity T-stems to overcome the thermodynamic penalty incurred by charge repulsion between their amino acid cargos and the negatively charged residues lining acceptor stem binding pocket of EF-Tu^102^. We then designed a new probe that targeted the altered T-stem, excluding the binding of wild-type 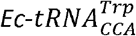. The mutant 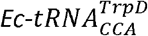 expressed well (**Supplemental Figure 7**) and was purified in high yield (10 nmol per binding) using our optimized purification method.

**Figure 7.**
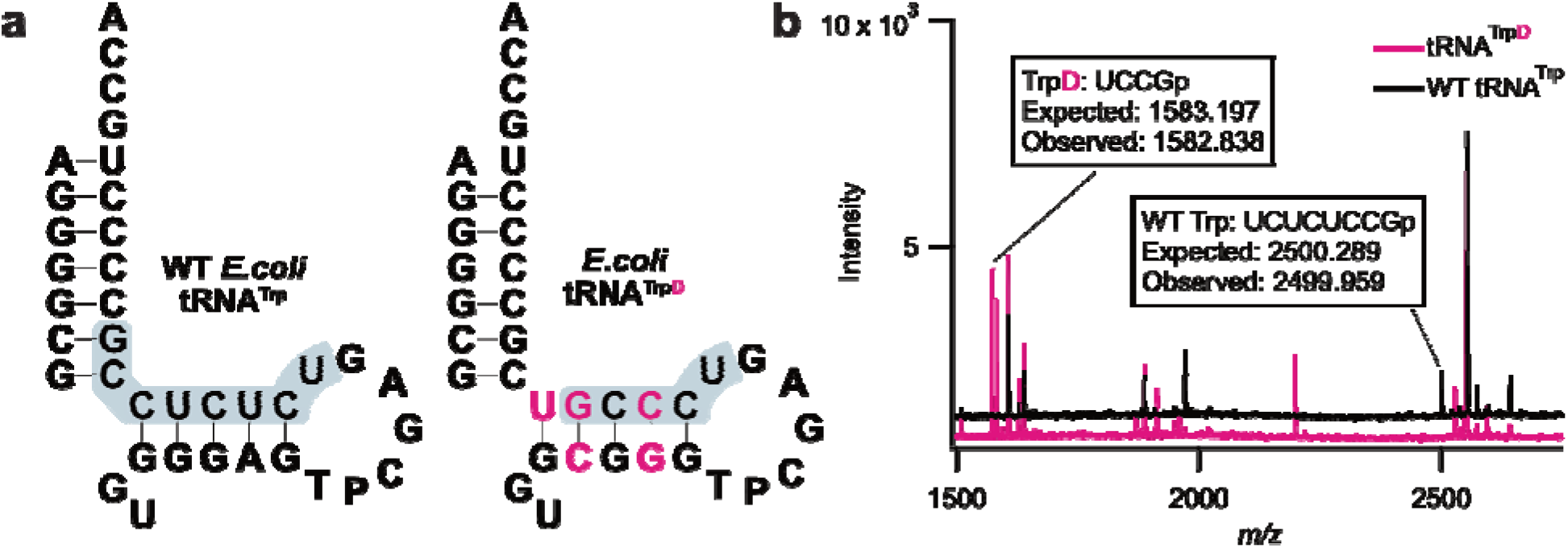
Mutant *Ec*-can be purified separate native *Ec*-. A) T-stems of wild-type *Ec*- and mutant *Ec*-. Sequences resulting from RNase T1 digestion are highlighted in grey. Bases mutated to match the *Ec*-T-stem are colored pink. B) MALDI spectra of wild-type (black trace) and mutant (pink trace) *Ec*-.

We employed both urea-PAGE and MALDI-TOF of RNAse T1 digested fragments to assess purity as before. PAGE revealed a single dominant band along with a faint band of higher molecular weight too large to be considered a contaminating endogenous tRNA (**Supplemental Figure 7**). MALDI-TOF revealed the, *Ec-*tRNA^*Asp*^T-stem (UCCGp, **Figure 7a**) along with other sequence elements unique to, 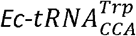. Importantly, the T-stem of wild-type tRNA^Trp^ (UCUCUCCGp, **Figure 7a**) was completely absent in the spectra (**Figure 7b**). Interestingly, 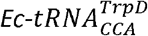 exhibited a greater heterogeneity in A37 modifications compared to wild-type 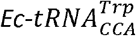 (**Supplemental Data File 1**), suggesting that the T-stem mutation may have interfered with the recognition of modification enzymes. This is surprising since the mutated bases are distal to the identity elements for MiaA recognition.

## Conclusions

In this study, we complemented previously described tRNA purification techniques with an overexpression strategy, enabling the purification of both native and non-native *E. coli* tRNAs. We produced tRNAs of great yield and purity, while maintaining native modifications. Our efforts revealed the modification profiles of tRNAs that are critical in the genetic code expansion efforts. Another key finding of our study also underscores the importance of post-transcriptional modifications in amber suppression. As further progress is made in both sense and stop codon reassignment, consideration should be given to the roles of post-transcriptional modifications in codon reassignment.

While the genes responsible for many post-transcriptional modifications have been known for years, the dynamics of the tRNA epitranscriptome are still being elucidated^103^. We identified a wide diversity of modification states for tRNAs unreported in the literature, reflecting this limited knowledge. Additional investigations will be required to understand the plurality modifications within tRNAs in response to growth conditions, stressors, nutrient abundances, and other factors. Considering the direct role post-transcriptional modifications play in the translation capacities of tRNAs, changes in modification stoichiometries could dramatically alter the proteome^104^. NGMS-seq can address these questions, offering direct quantitation of modification stoichiometries rather than inferring their abundances using conventional sequencing approaches^105^.

tRNA is of crucial importance in PURE CFES, but no commercial source of fully modified tRNA currently exists (Roche MRE600 tRNA reagent was discontinued in 2021). This work demonstrates a solution to that problem, providing a method for overexpression and purification of natively modified *E. coli* tRNAs. We anticipate that our approach we present will allow researchers to build custom tRNA pools, potentially maximizing the expression yields of PURE CFES^104^and enabling more robust genetic code reassignment^32^, as well as broader applications that may benefit from pure, fully modified tRNAs.

## Acknowledgments

E.M.K., J.L.A., L.D-F., J.A.D. and K.P.A were supported by Alfred P. Sloan Foundation grant G-2024-22710 and by NSF Award 2419641. Intact tRNA LC-MS and HPLC purification of tRNA was supported by the NSF Center for Genetically Encoded Materials (CHE-2002182; I.K., E.M., A.S., K.C. and S.C.B.). Grant support for construction of the 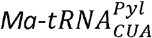 expression plasmid was provided by the National Institutes of Health (5R01GM079238 to S.C.B.). The authors also acknowledge grant support from the US National Institutes of Health R01 HG012853, R41 HG013624, R41 HG014125, U24 HG011735, and RM1 HG011563 (to S.Z), and NSF award 2338121 and US National Institutes of Health R01 GM152459 (to A.E.E). We separately thank Prof. Ya-Ming Hou and Prof. Anna Kashina for the gift of pKK223-3_ArgII.

## Materials and methods

### Creation of tRNA expression vectors

Plasmids for the expression of each tRNA were generated through Gibson assembly. Briefly, the pKK223-3 backbone was linearized using Q5 PCR (New England Biolabs, Ipswich, MA, USA). Gibson assembly inserts generated from overlapping PCR from primers designed using Primerize^107,108^. 20 µL Gibson assembly reactions used a 7:1 ratio of insert to backbone and were incubated for 1 hr at 50°C. The plasmid bearing the pUC19 origin of replication was similarly created by Gibson assembly of 4 fully synthetic DNAs. Assembled constructs were transformed into chemically competent DH5α (New England Biolabs, Ipswich, MA, USA) cells using heat-shock and selected using antibiotic selection plates containing carbenicillin at a final concentration of 100 µg/mL. Successful transformants were sequenced using whole plasmid sequencing from Plasmidsaurus (San Francisco, CA).

### Overexpression of tRNAs and isolation of enriched tRNA pools

Isolation of enriched tRNA pools was performed according to Avcilar-Kucukgoze^1^. In summary, for each tRNA, 1L of LB broth containing a final concentration 100 µg/mL carbenicillin was inoculated with a 10 mL overnight culture. Cultures were grown at 37°C in a shaking incubator at 250 rpm to an OD_600_ of 0.5 and then induced with 300 µM IPTG. Cultures were allowed to induce overnight (~16hrs) at 37°C before harvesting by centrifuging at 4,500xg at 4°C for 30 minutes. The resulting cell pellets were washed once with 30 mL of 0.9% NaCl before storage −80°C until processing.

For the generation of an enriched 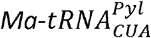 pool, pUC19 vector containing 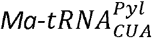 under the control of a Pm promoter and *Ma*PylRS under the control of arabinose was transformed into DH5α cells. Overnight cultures were used to inoculate 1 L of Terrific Broth containing 100 µg/mL and grown to an OD600 of 0.6 at 37°C in a shaking incubator. Once OD was achieved, both tRNA and AARS expression was induced with the addition of toluic acid (f.c. 300 µM) and arabinose (f.c. 0.01% w/v); 1 mM Boc-lysine (Chem-Impex) was also added to charge the tRNA. The culture was allowed to induce overnight at 25°C and harvested by centrifugation of 4,500x g at 4°C for 30 minutes. The cell pellet was washed once with 30 mL of 0.9% NaCl before storage at −80°C until processing.

To recover the enriched tRNA pools, cell pellets were resuspended in 18 mLs of Extraction Buffer (50 mM sodium acetate, 10 mM MgOAc pH 5.0). Nucleic acids were extracted by adding 17.2 mL of acidic phenol pH 4 (VWR International, Radnor, PA, USA) and were shaken at room temperature using a shaking incubator for 30 min. The resulting emulsion was partitioned into aqueous and organic phases by centrifugation at 4,500x g for 15 min at 4°C. The aqueous phase was collected and the organic phase extracted once more by adding 14 mL of Extraction Buffer and shaking for 15 min followed by another round of centrifugation. Aqueous phases were pooled together, and the total nucleic acid fraction was precipitated by the addition of 1.5 mLs of 5M NaCl followed by the addition of 1 volume of isopropanol and centrifugation at 14,500x g for 15 min at room temperature. The pellet was washed briefly with ice-cold 70% ethanol before briefly drying. Next, the pellet was resuspended in 15 mL of cold 1 M NaCl and centrifuged at 9,500xg for 20 min at 4°C. The supernatant was collected before being precipitated with 30 mLs of ice-cold 100% ethanol by incubating for at least 30 minutes at −20°C. The nucleic acids were recovered by centrifuging at 14,500xg for 5 min at 4°C. The resulting pellet was washed briefly with ice-cold 70% ethanol and resuspended by gentle mixing of 6 mL of 0.3 M NaOAc, pH 5.0. The remaining DNA was precipitated by adding 3.4 mL isopropanol and incubating for 10 minutes at room temperature before centrifugation at 14,500x g for 5 min at room temperature. The supernatant was collected and the tRNAs were precipitated by adding 2.3 mL of isopropanol and incubating at −20°C overnight. The precipitated tRNA pool was centrifuged at 14,500xg for 15 min at 4°C, washed once with ice cold 70% ethanol, and allowed to air dry. Finally, the tRNA pellet was resuspended in 500 µL of ddH_2_O and quantified using a Nanodrop ND-1000 spectrophotometer (Thermo Fisher Scientific, Waltham, MA, USA).

### Coupling of hybridizing probes for Native E. coli tRNA purification

Probes for the purification of native *E. coli* tRNAs were taken from Miyauchi et al.^40^ and synthesized from Integrated DNA Technologies (Coralville, IA, USA) containing a /3AmMO/ group for NHS coupling. For each tRNA, probe couplings to premade HiTrap-NHS FF columns (Cytiva Life Sciences, Marlborough, MA, USA) were performed according to the manufacturer’s protocol. Briefly, lyophilized probes were first resuspended to a final concentration of 1 mM in ddH2O and diluted to concentration of ~4 A_260_ units in Coupling Buffer (0.2 M NaHCO_3_, 0.5 M NaCl, pH 8.3) to a volume of 1 mL. To prepare columns for coupling, the column storage buffer was displaced and removed by injecting 6 mLs of ice-cold 1 mM HCl into the column using a syringe. The column was then loaded with 1 mL of the previously prepared probe solution using a syringe and allowed to couple to the resin for 30 minutes at room temperature. Following the coupling reaction, 3 mLs of Coupling Buffer was used to flush the column of unreacted probe and 1 mL of flow-through was collected to quantify the yield of the coupling reaction. Unreacted resin was blocked by flushing the column first with 6 mLs of Blocking Buffer A (0.5 M ethanolamine-HCl, 0.5 M NaCl, pH 8.3), 6 mLs of Blocking Buffer B (0.1 M NaOAc, 0.5 M NaCl, pH 4.0), and a final flush of 6 mLs Blocking Buffer A. Blocking was allowed to occur for 30 minutes at room temperature. Following the blocking reaction, the column was flushed again by washing with 6 mLs Blocking Buffer B, 6 mLs of Blocking Buffer A, and finally 6 mLs of Blocking Buffer B. Columns were then prepared for storage by washing with 6 mLs of Column Storage Buffer (50% ethanol, 50 mM NaOAc, pH 5.0) and kept at 4°C until use. To determine the coupling yield and estimated binding capacity of the column, the 1 mL of previously recovered flow-through was acid-neutralized with 1 mL of 2M glycine pH 2.0 and A_260_ was measured using a Nanodrop spectrophotometer.

### Purification of individual tRNAs from overexpressed tRNA isolate

The purification of individual tRNAs was adapted from the protocol described by Miyauchi et al^40^. First, 100 µL aliquot of isolated, enriched tRNA pool was diluted into 10 mL of tRNA Binding Buffer (0.9 M tetramethylammonium-HCl, 0.1 mM EDTA, 10 mM Tris-HCl pH 7.4) and kept on ice until binding. Purifications were performed using an ÄKTA start FPLC, though any peristaltic pump system should suffice. The oligonucleotide probe column for the tRNA of interest was first attached to the ÄKTA start and equilibrated by running 10 mL of tRNA Binding Buffer using a flow rate of 1 mL/min. The flow rate of 1 mL/min was maintained for the duration of the purification. Following equilibration, the column was placed in a water bath set to 75°C and allowed to heat for 10 minutes. After warming the column, 10 mL of enriched tRNA pool were washed over the column in a recirculating manner where the inlet and outlet tubes drew from and fed into the same reservoir. The temperature of the water bath was slowly cooled to 60°C (~30 min) and the tRNA was allowed to bind at this temperature for at least 10 minutes. After binding, the ÄKTA was paused, and the column was removed from the water bath to cool briefly at room temperature. Unbound tRNAs were washed from the column by running at least 15 mLs of Wash Buffer (10 mM Tris-HCl pH 7.4). Washing was monitored by measuring the A_260_ values of flow-through via Nanodrop and was completed after A_260_ was <0.01 units. After washing, the column was removed from the ÄKTA and capped before a syringe filled with 4 mLs of wash buffer was attached using a luer-lock adapter. To elute, the column-syringe apparatus was submerged in a water bath set 80°C and incubated for 5 minutes. Immediately after the 5-minute incubation, the heat-eluted tRNAs were collected by flushing the column with 1.1 mLs of Wash Buffer using the pre-attached syringe and immediately cooled on ice. A second elution step was performed once more to ensure no tRNAs remained on the column. The yield of each elution step was estimated using a Nanodrop. The binding, wash, and elution steps can be done several times depending on the binding capacity of the column. For the tRNAs in this work, two bindings were performed per 100 µL aliquot, unless otherwise specified. Eluted fractions were pooled together and precipitated overnight at −20°C by adding 0.1 volume of 3M NaOAc pH 5.3 and 1 volume of isopropanol. After precipitation, tRNAs were centrifuged for 10 minutes at 4C at 16,000xg and washed once with ice-cold 70% ethanol. Precipitated tRNAs were resuspended in ddH2O and concentration was quantified via Nanodrop.

### Cell-free expression of HiBiT with HiBiT-specific tRNAs

The ten purified tRNAs required for HiBiT translation were first individually refolded in RNase-free water by heating to 95°C and stepwise cooling to 25°C over 10 minutes. The refolded tRNAs were then pooled together in equal concentrations compromising a tRNA pool with a pooled concentration of 2 mg/mL. Cell-free expression was carried out in NEB PURExpress (ΔtRNA,ΔAA) (New England Biolabs, Ipswich, MA, USA) in 12.5 µL reactions where the tRNA pool was supplied by either the custom pool or MRE600 (Roche, discontinued), both at a final concentration of 0.2 mg/mL. A tRNA-free reaction served as the negative control. The HiBiT DNA template was generated using Primerize and added at a final concentration of 10 nM. Translation reactions were carried out at 37°C for 2 hrs in a thermocycler. HiBiT translation yields were quantified by diluting the PURE reactions 100-fold in ddH_2_O. Then, 10 µL of each dilution was reacted with 10 µL of 1 X NanoGlo Extracellular reagent (Promega, Madison, WI, USA) within the wells of a white 384-well plate. Luminescence was tracked for 20 minutes and monitored for a peak, steady-state luminescence signal using a SpectraMAX Gemini EM MicroPlate Reader.

### Design of 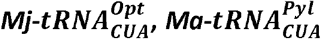, and 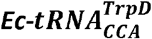 hybridization probes

The purification probes for non-native tRNAs were designed according to principles described by Yokogawa et al^109^. Oligonucleotides for 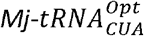 and 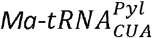 purifications were first designed to cover roughly 30 nts of each tRNA spanning from anticodon to the T-stem of the tRNA to achieve a T_M_ of ~65°C. Then, candidate probes were checked for potential homodimer and monomer secondary structure formation and altered to reduce their potential formations. In a similar fashion, 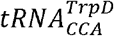 probe was designed to span both the 3’-tail and mutant T-stem required for selective purification against wild-type 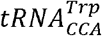.Finally, each probe was designed to include an 3’-AmmO group for coupling to the NHS resin.

The 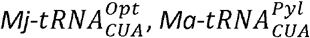, and 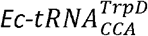 probes were coupled in a similar fashion to the native *E. coli* tRNA probes with the exception that they were coupled with ~20 A_260_ units of probe to facilitate higher yields of purification.

### Preparation of *in vitro* transcribed 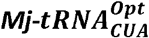 and 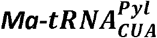

DNA template sequence for *in vitro* transcription of 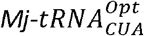 transzyme were taken from Martin et. al^110^ and assembled using Primerize-designed overlapping PCR. Product was confirmed on a 3% agarose gel before being purified using the Epoch PCR Clean-up Kit (Epoch Life Sciences, Inc., Missouri City, TX, USA). The purified PCR product was then used in a 200 µL *in vitro* transcription reaction (0.2 μM template, 1X Homemade NEB Transcription Buffer, 8 mM GTP, 4 mM A/C/UTP, inorganic pyrophosphatase 25 ng/μL, 1 μM T7 RNAP, Murine RNAse Inhibitor (New England Biolabs, Ipswich, MA, USA) 0.4 U/μL. The assembled reaction was allowed to react 16hrs at 37°C. Following transcription, 4 units of TubroDNAse (Thermo Fisher, Waltham, MA, USA) were added to the reaction and incubated for 30 min at 37°C degrade the DNA template.

After degradation of the DNA template, hammerhead-cleavage was facilitated by diluting the reaction 5-fold in a solution of 30 mM MgCl_2_ and 0.2 mM EDTA and incubating at 60°C for 2hrs. The reaction was subject to phenol:chloroform extraction and ethanol precipitation overnight using glycogen as a carrier. The cleaved 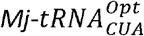 product was then gel-purified from a 10% (19:1) polyacrylamide gel and again ethanol precipitated overnight using glycogen as a carrier. Precipitated tRNA was resuspended in ddH_2_O and concentration was quantified using a Nanodrop spectrophotometer.

### Preparation of *in vitro* transcribed 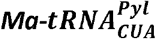

*In vitro* 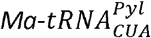 was prepared using *in vitro* transcription. DNA template was first assembled using PCR from Primerize designed oligonucleotides synthesized by IDT. *In vitro* transcription reactions were assembled as above in a 200 uL reaction volume: 0.2 μM template, 1X Homemade NEB Transcription Buffer, 8 mM GTP, 4 mM A/C/UTP, inorganic pyrophosphatase 25 ng/μL, 1 μM T7 RNAP, Murine RNAse Inhibitor (New England Biolabs, Ipswich, MA, USA) 0.4 U/μL. Assembled *in vitro* transcription reaction were allowed to react for 8 hrs at 37°C. Following the reaction, DNA template was digested using 4 units of TURBO DNAse for 30 minutes at 37°C. tRNA was purified using a Monarch Spin RNA Clean-up Kit (New England Biolabs, Ipswich, MA, USA) and eluted into nuclease-free, ddH_2_O. Aliquots of purified tRNA were flash frozen and stored at −80°C.

### Cell-free translation of sfGFPT216TAG and fMet-Gly-TAG-HiBiT-FLAG

Amber suppression CFES was carried out using PURExpress (New England Biolabs, Ipswich, MA, USA). 30 µL reactions containing 10 µM pCNFRS, 1 mM L-paracyanophenylalanine (Chem-Impex International, Wood Dale, IL, USA), 5 µM of either *in vivo* or *in vitro* 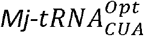, and 10 nM of either the fMet-Gly-TAG-HiBiT-FLAG or sfGFPT216TAG templates were assembled and incubated for 2 hrs at 37°C. Reactions containing no 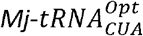 served as a negative control. For sfGFPT216TAG, amber-codon suppression yield was determined by measuring the emission at 509 nm from excitation at 488 nm. For fMet-Gly-TAG-HiBiT-FLAG constructs, a 2 µL aliquot of each reaction was first diluted 100-fold into ddH_2_O. Then, 10 µL of each dilution was reacted with 10 µL of 1X NanoGlo Extracellular reagent (Promega, Madison, WI, USA). Luminescence was tracked for 20 minutes and monitored for a maximal, steady-state luminescence signal.

### MALDI-TOF mass spectrometry of PURE CFES translated fMet-Gly-HiBiT-FLAG

The translated fMet-Gly-TAG-HiBiT-FLAG peptides were purified using magnetic bead, FLAG-tag purification using a magnetic bead separation plate according to manufacturer’s instructions. For each reaction, 20 µL of well mixed Anti-FLAG M2 Magnetic Beads (Millipore Sigma, Burlington, MA, USA) were aliquoted into 1.5 mL centrifuge tubes. Beads were equilibrated 5 times with phosphate buffered saline (PBS: 137 mM NaCl, 2.7 mM KCl, 10 mM Na_2_PO_4_, 1.8 mM KHPO_4_) before final resuspension with 20 µL of PBS. The remaining volume (25 µL) of PURE CFES reactions translating fMet-Gly-TAG-HiBiT-FLAG were diluted into 400 µL of PBS and were added to resuspended beads. Diluted reactions were allowed to bind to the beads using end over end mixing for 30 minutes at room temperature.

Following binding, beads were collected using magnetic separation and the flow-through was discarded. Beads were subsequently washed 5 times with volume of PBS and were eluted by the incubation of 0.1% trifluoroacetic acid for 15 minutes and removal from the beads.

Eluted peptides were then prepared for MALDI by first ZipTip desalting using a C_18_ stationary phase. Desalted samples were then spotted on the MALDI target on top of a solution of saturated alpha-cyano-4-hydroxycinnamic acid matrix dissolved in 50:50 acetonitrile in water. Peptides were analyzed using a Bruker Autoflex Max MALDI/TOF-MS (Bruker, Billerica, MA, USA) in reflectron positive mode. Resulting spectra were analyzed in the mMass openware software (https://github.com/xxao/mMass) first by smoothing spectra with 2 cycles of Savitzky-Golay smoothing over a window of 0.2 *m/z* using the Peak Picker algorithm. Peaks were filtered using S/N ratio >3.0 and relative peak intensity of >0.5%.

### Profiling of tRNA RNAse T1 and Fragments using MALDI-TOF

Sequence-specific digestion products of purified tRNAs were analyzed following a protocol established by Hossain and Limbach with minor modifications^45^. Roughly 1-2 µg of each purified tRNA were diluted into 20 µL of 200 mM NH_4_OAc pH 5.2, heat denatured at 95°C for 2 minutes and cooled immediately on ice. After a brief incubation on ice, 50 units of RNAse T1 (Thermo Fisher, Waltham, MA, USA)per microgram of tRNA were added and allowed to incubate at 37°C for 30 minutes. Digested fragments were analyzed using Brucker Autoflex MALDI-TOF in reflectron negative mode by first pre-spotting targets with 0.5 µL of 300 mM trihydroxyacetophenone (THAP) in neat acetonitrile (ACN). Samples were mixed with an equal volume of 1:1 300 mM THAP in ACN and 250 mM diammonium hydrogen citrate in 100% ddH_2_O. 1 µL of prepared sample was then overlaid onto pre-spotted THAP. Following data acquisition, spectra were smoothed using 2 cycles of Savitzky-Golay and a window size of 0.2 *m/z* using the mMass Software. Peaks were assigned using the mMas Peak Picker tool with filtering peaks to a S/N of 3.0 and relative signal intensity of >0.5%. Expected RNAse T1 digestion products were generated *in silico* using the MongoOligonucleotide Calculator (http://mass.rega.kuleuven.be/mass/mongo.htm) using literature reported modified tRNA sequences obtained from MODOMICS (https://genesilico.pl/modomics/). Spectra were analyzed for constituent tRNA fragments of each purified tRNA, potential hypomodified variants of these fragments, evidence of isoacceptor sequences, and signature digestion products of all *E. coli* tRNA sequences.

### Intact tRNA LC-MS of native E. *coli* tRNAs

Intact tRNA LC-MS was performed as previously reported^46^. Samples were resuspended in water before analysis using intact tRNA LC-MS. In general, 20 pmol of tRNA was injected to ensure adequate signal in the UV and ion chromatograms. Samples were resolved on a ACQUITY UPLC BEH C18 Column (130 Å, 1.7 μm, 2.1 mm X 50 mm, Waters part # 186002350, 60 °C) using an ACQUITY UPLC I-Class PLUS (Waters part # 186015082). The mobile phases used were (A) 8 mM TEA, 80 mM HFIP, 5 μM EDTA (free acid) in 100% MilliQ water; and (B) 4 mM TEA, 40 mM HFIP, 5 μM EDTA (free acid) in 50% MilliQ water/50% methanol. Samples were eluted using a flow rate of 0.3 mL/min. The elution method began with Mobile Phase B at 22%; the fraction of Mobile Phase B increased linearly to 40% B over 10 min and from 40 to 60% B over 1 min and was then held at 60% B for 1 min. The fraction of Mobile Phase B was then decreased linearly from 60 to 22% B over 0.1 min and held at 22% B for 2.9 min to equilibrate the column. For more difficult separations, the method was extended as needed. The mass of the RNA was analyzed using LC-MS with a Waters Xevo G2-XS Tof (Waters part #186010532) in negative ion mode with the following parameters: capillary voltage: 2000 V, sampling cone: 40, source offset: 40, source temperature: 140 °C, desolvation temperature: 20 °C, cone gas flow: 10 L/h, desolvation gas flow: 800 L/h, 1 spectrum/s. Deconvoluted masses were obtained using the MaxEnt software (Waters Corporation).

### Peak assignments of native *E. coli* tRNAs analyzed by Intact tRNA LC-MS

Mass spectra were analyzed using Mmass’s Peak Picker algorithm selecting for peaks with a S/N of 10 and relative threshold of 5%. Native *E. coli* sequences complete with post-transcriptional modifications were taken from MODOMICS and masses were calculated using MongoOligonucleotide Calculator. For tRNAs expected to contain the ct6A modification, a custom user base anion with mass of N=261.217 *m/z* was used. Masses were then assigned first according to literature reported sequences before considering potential hypomodifications or additional modifications. Peak areas of tRNA species were calculated using a custom macro implemented in the Igor Pro software.

### Chemi-luminescence Northern Blot

To detect 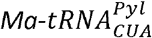 expression, we performed chemiluminescent northern blotting^111^. Total tRNA isolates from 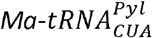-expressing cells, along with an *in vitro* transcribed 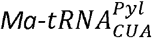 (positive control) and *E. coli* MRE600 tRNA (negative control), were separated by denaturing urea-PAGE and transferred to a positively charged nylon membrane (Immobilon-Ny+, Millipore Sigma, Burlington, MA, USA) via electroblotting and UV-crosslinked. Crosslinked membranes were pre-hybridized in Ultrahyb Ultrasensitive buffer (Thermo Fisher, Waltham, MA, USA), then hybridized overnight at 42°C with a 51-biotinylated DNA probe (5 nM final concentration).

Following hybridization, membranes were washed twice with 2× SSC and 0.1% SDS, and twice with 0.1× SSC and 0.1% SDS at 42°C. Detection was carried out using the Chemiluminescent Nucleic Acid Detection Module (Thermo Fisher, Waltham, MA, USA) according to the manufacturer’s protocol. Blots were blocked for 15 minutes with gentle rocking, incubated with 50 µL streptavidin-HRP for 15 min, and washed five times with the supplied wash buffer. After equilibration, chemiluminescent substrate (luminol/enhancer and peroxide) was applied, and the resulting signal was captured using automatic exposure on an OMEGA LUM™ G Imaging System (Aplegen, Gel Company, Inc, San Francisco, CA).

### Separation of 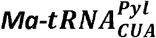 using Hydrophobic Interaction Chromatography

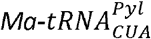 fractions were separated from tRNA bulk previously obtained by DNA probe isolation using a TSK Phenyl-5PW hydrophobic interaction chromatography (HIC) column (Tosoh Bioscience, Tokyo, Japan) with a linear gradient starting from buffer A (10 mM NH_4_OAc, pH = 5.8; 1.7 M (NH_4_)_2_SO_4_) to buffer B (10 mM NH_4_OAc, pH = 5.8; 10% MeOH). Elution fractions corresponding to F3 (27 min) were pooled, dialyzed into MiliQ water, and concentrated using Amicon ultra centrifugal filter (10 kDa MWCO) (Millipore Sigma, Burlington, MA, USA).

### NGMS Sequencing of 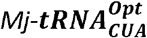 and 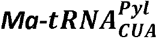

tRNA samples for NGMS-Seq were split into two portions. 10% (0.1-1 pmol) were mixed with RNase-free water and analyzed by LC–MS without acid hydrolysis. The remaining samples underwent formic acid hydrolysis to generate RNA ladder fragments for MS sequencing. Controlled hydrolysis of tRNA, LC-MS OE240 analysis, NGMS-Seq data analysis, and stoichiometry determination followed previously published protocols^70,112,72,71^ with minor adjustments described below.

### Controlled Hydrolysis of tRNA

tRNA samples (1-15 pmol) were first mixed with 10⍰µL of RNase-free water and transferred to 0.5⍰mL Eppendorf tubes. The samples were denatured at 70⍰°C for 15 minutes using a thermal cycler. To initiate controlled acid hydrolysis, an equal volume of 1% formic acid was added to the denatured sample. The mixtures were incubated at 70⍰°C for 5 minutes, after which the tubes are immediately quenched on dry ice for 5 minutes. Then samples were dried to completion using a Vacufuge Plus concentrator (Eppendorf, Enfield, CT, USA). The resulting dried residues were then reconstituted in RNase-free water for LC–MS measurement.

### LC-MS OE240 measurement

A DNAPac RP column (2.1*50mm DNAPac RP) was purchased from Thermo Fisher (Waltham, MA, USA) and installed in a Vanqiush Horizon UHPLC system (Thermo Fisher, Waltham, MA, USA). Mobile phase A was prepared with 15 mM diisopropylamine (DIPA) and 35 mM 1,1,1,3,3,3-hexafluoroisopropanol (HFIP) buffers in aqueous solution. Mobile phase B was prepared at the ratio of 50:50 water/methanol with buffer 15 mM DIPA and 35 mM HFIP.

#### LC conditions

Column temperature was maintained at 60 °C and the flow rate was set at 0.25 mL/min. The gradient for mobile phase B was as follows: 1% at 0 to 1⍰min, increased linearly to 45% over 21⍰min, ramped to 95% from 21 to 23⍰min, and returned to 1% by 23.5⍰min.

#### MS conditions

tRNA samples were analyzed in negative-ion full MS mode from m/z 600 to 2000 with a scan rate of 2 spectra/s at 120k resolution. Sheath gas was set at 45 Arb and aux gas was set at 10 Arb. Ion transfer tube temperature was set at 320°C and the vaporizer temperature was set at 275°C.

### NGMS-Seq data analysis

The experimental output data from Biopharma Finder 5.0 was exported as .xlsx files. Sequencing was performed using a similar method described in the references^70–72^.

### Partial RNA modification stoichiometry determination

Each partial nucleotide modification creates a branch in the 2D mass-t_R_ ladder that is evident in all subsequent positions in the 2D sequence ladder, so this stoichiometry calculation was repeated in multiple post branch positions for each partial modification to obtain six values that were used to calculate averages and standard deviations unless indicated otherwise.

For example, relative MS intensity ratios of ladder fragments containing mG and canonical G at post branch position 46 were used to calculate their stoichiometry at this position. This calculation was repeated in multiple subsequent post branch positions for the partial modification to obtain at least six values that were used to calculate averages and standard deviations

## Supplementary Information

**Figure S1.**
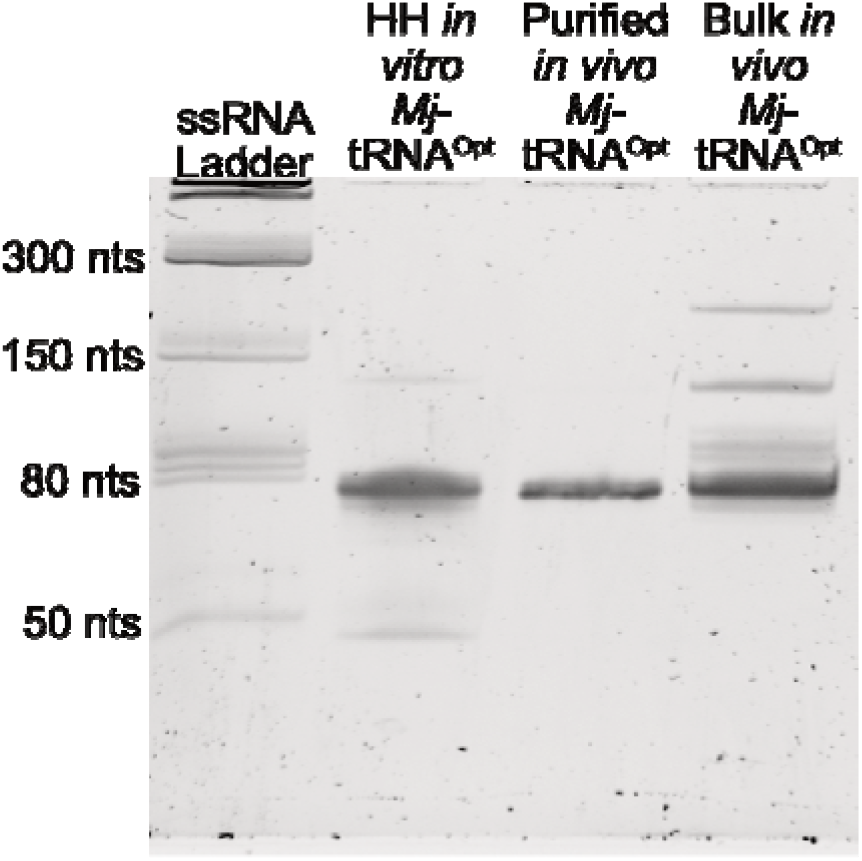
*Mj*-is greatly overexpressed and highly pure. Urea-PAGE gel of *Mj*-products. Lanes from left to right: ssRNA ladder, Hammerhead ribozyme (HH) cleaved and gel-purified in vitro *Mj*-transzyme, purified *in vivo Mj*-, and bulk overexpressed *in vivo Mj*-.

**Figure S2.**
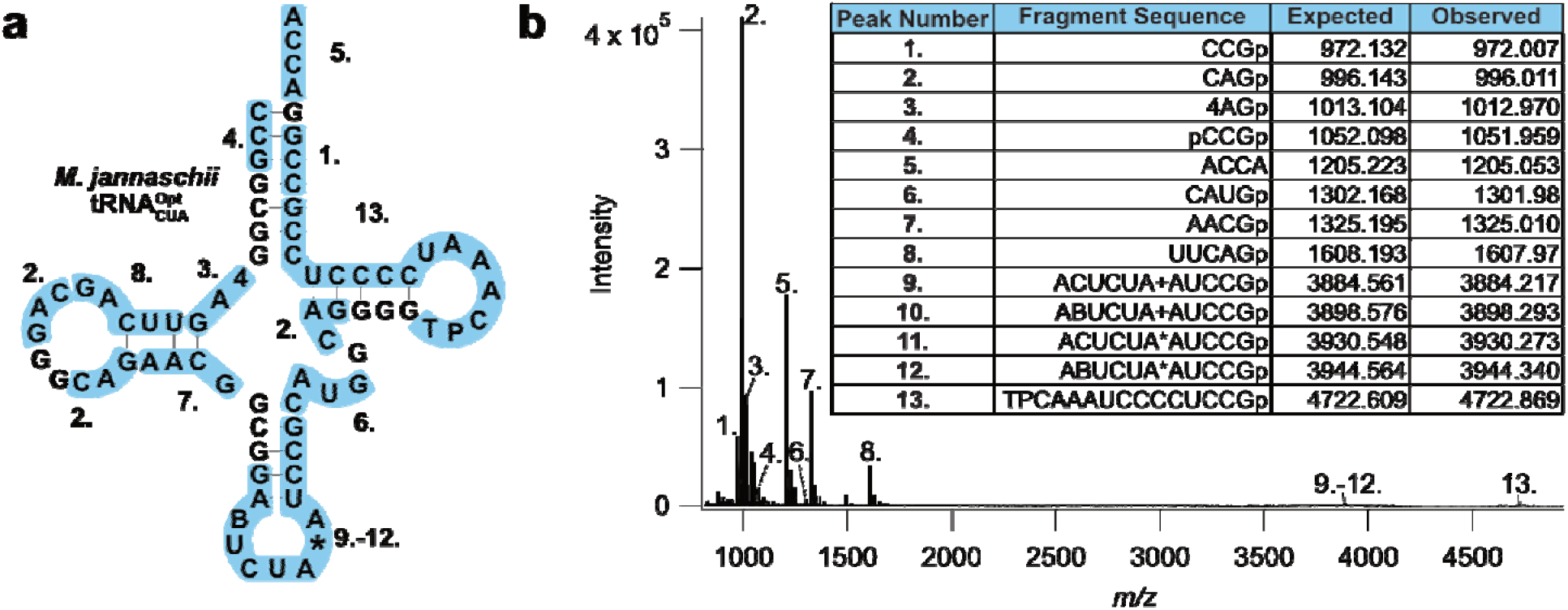
Draft modification profile of *Mj*-. **(a)** Draft modification profile of *Mj*-reconstructed from (b) RNAse TI fingerprinting of purified *in vivo Mj*-analyzed by MALDI-TOF. *Ma*sses and expected sequences are numbered in increasing expected masses.

**Figure S3.**
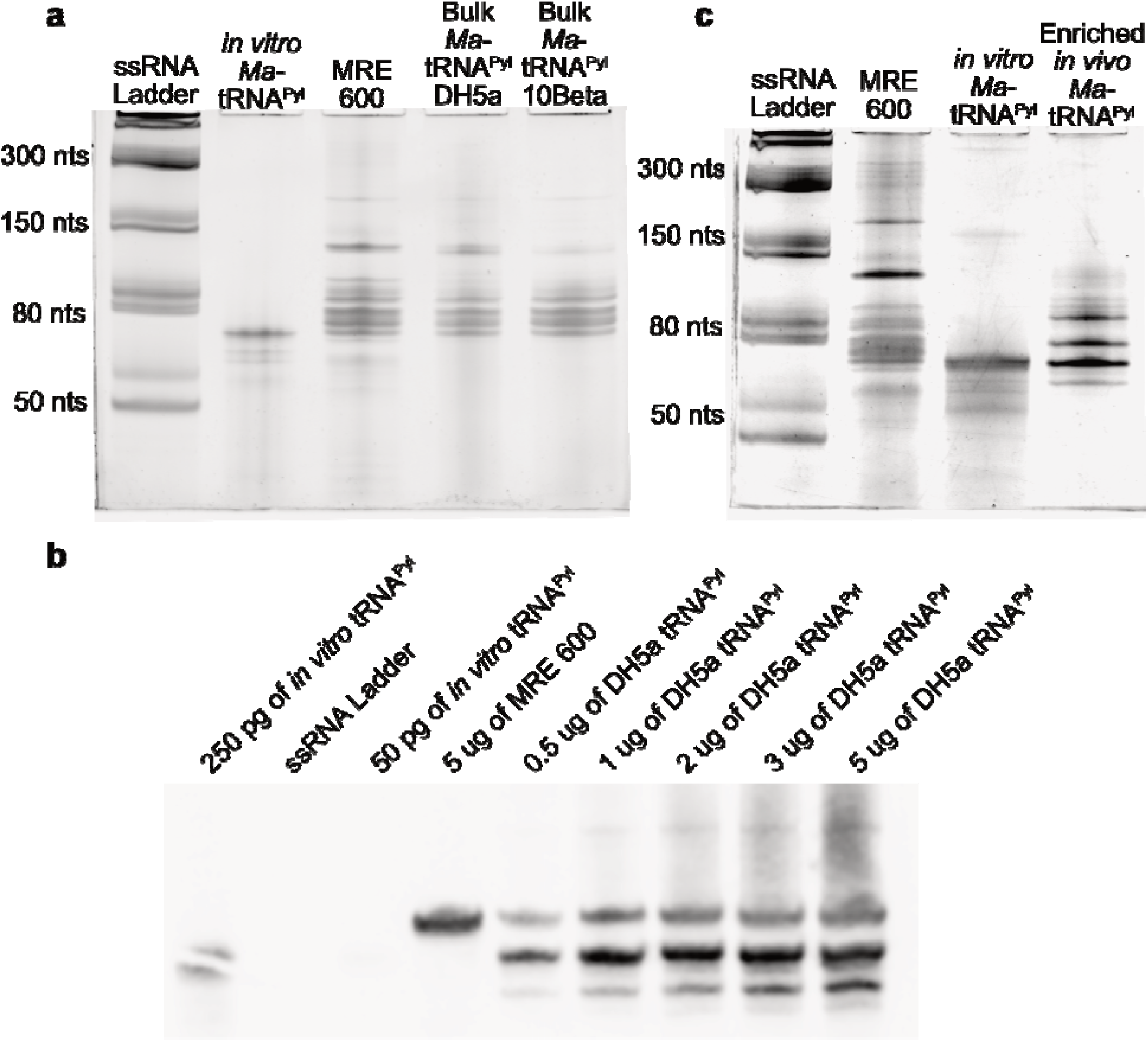
*Ma-* is poorly expressed in pKK223-3 and recalcitrant to purification. **(a)** Urea-PAGE gel of *Ma*-expressed in either NEB DH5alpha or NEB10beta cells. In vitro transcribed *Ma-* was used as a positive control. MRE600 tRNA served as a negative control. Gel was stained with SYBR Gold. **(b)** Chemi-luminescent Northern Blot of *Ma*-expressed in NEB DH5alpha. Either 250 pg or 50 pg of in vitro *Ma*-served as a positive control. 250 pg sample of in vitro *Ma*-tRNAPyl was nicked. Single-stranded RNA ladder and MRE600 served as negative controls. Isolated small RNA fractions of DH5alpha expressing 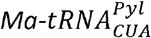 were loaded in increasing amounts.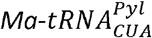 is indeed expressed though poorly in DH5α. A native *E. coli*tRNA non-specifically binds to the Northern probe designed for Ma-tRNAPyl but exhibits a higher molecular weight than the 71 nt long 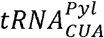**(c)** Urea-PAGE gel of 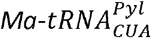 enriched through purification. Several native *E. coli* tRNA were co-purified with 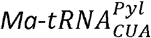.

**Figure S4.**
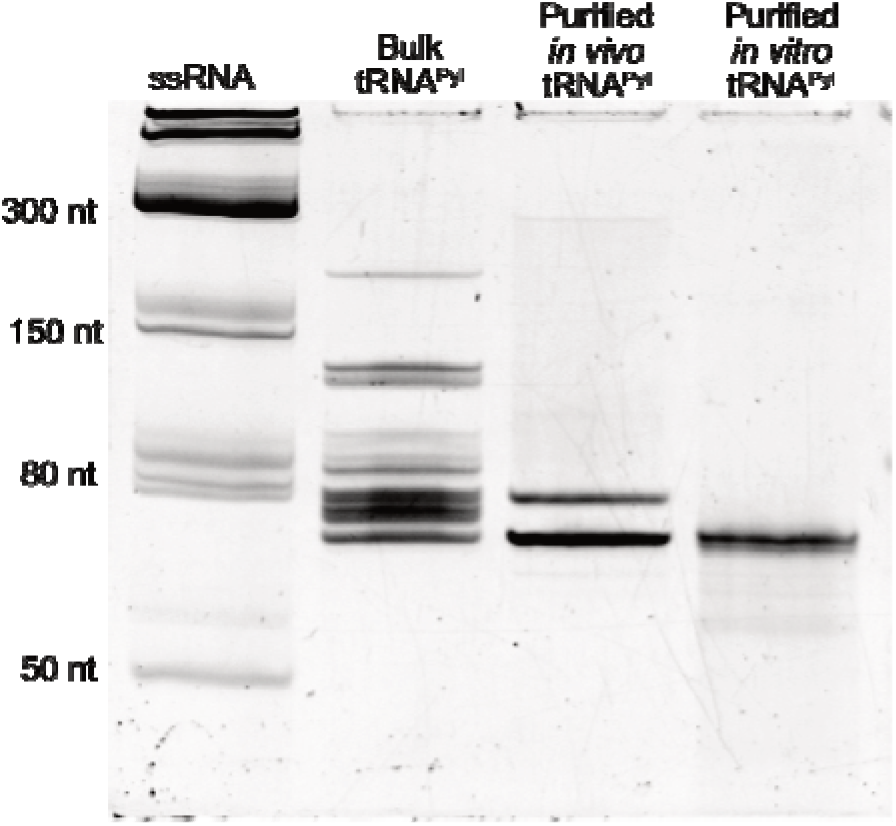
*Ma*-is greatly overexpressed and highly pure when expressed from the Pm:XylS controlled pUC19 vector. Urea-PAGE gel of overexpressed *Ma*-. Land assignments from left to right. Lane 1: ssRNA ladder, Lane 2: Bulk expressed and isolated *Ma*-, Lane 3: Purified *in vivo Ma*-from pUC19 expressed DH5alpha. Lane 4: in vitro *Ma*-.

**Figure S5.**
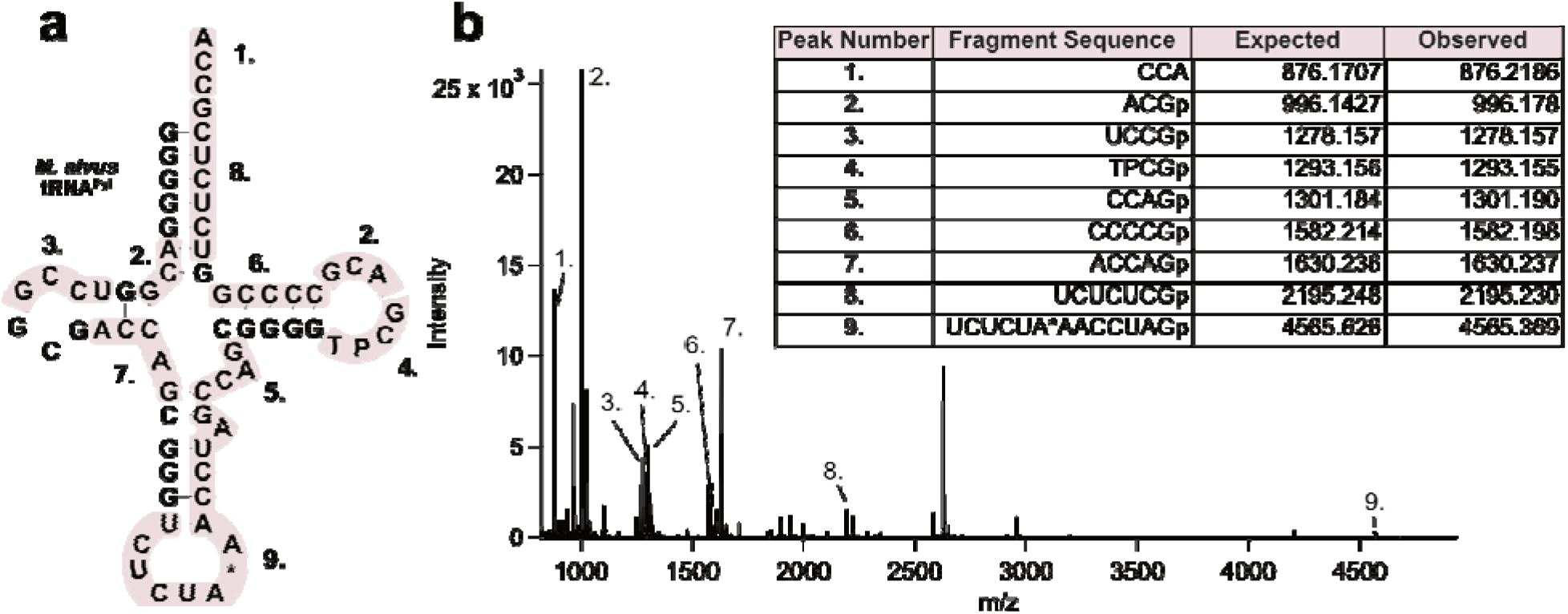
Draft modification profile of *Ma*-. **(a)** Draft modification profile of *Ma*-reconstructed from **(b)** RNAse TI fingerprinting of purified *in vivo Ma*-analyzed by MALDI-TOF. *Ma*sses and expected sequences are numbered in increasing expected masses.

**Figure S6.**
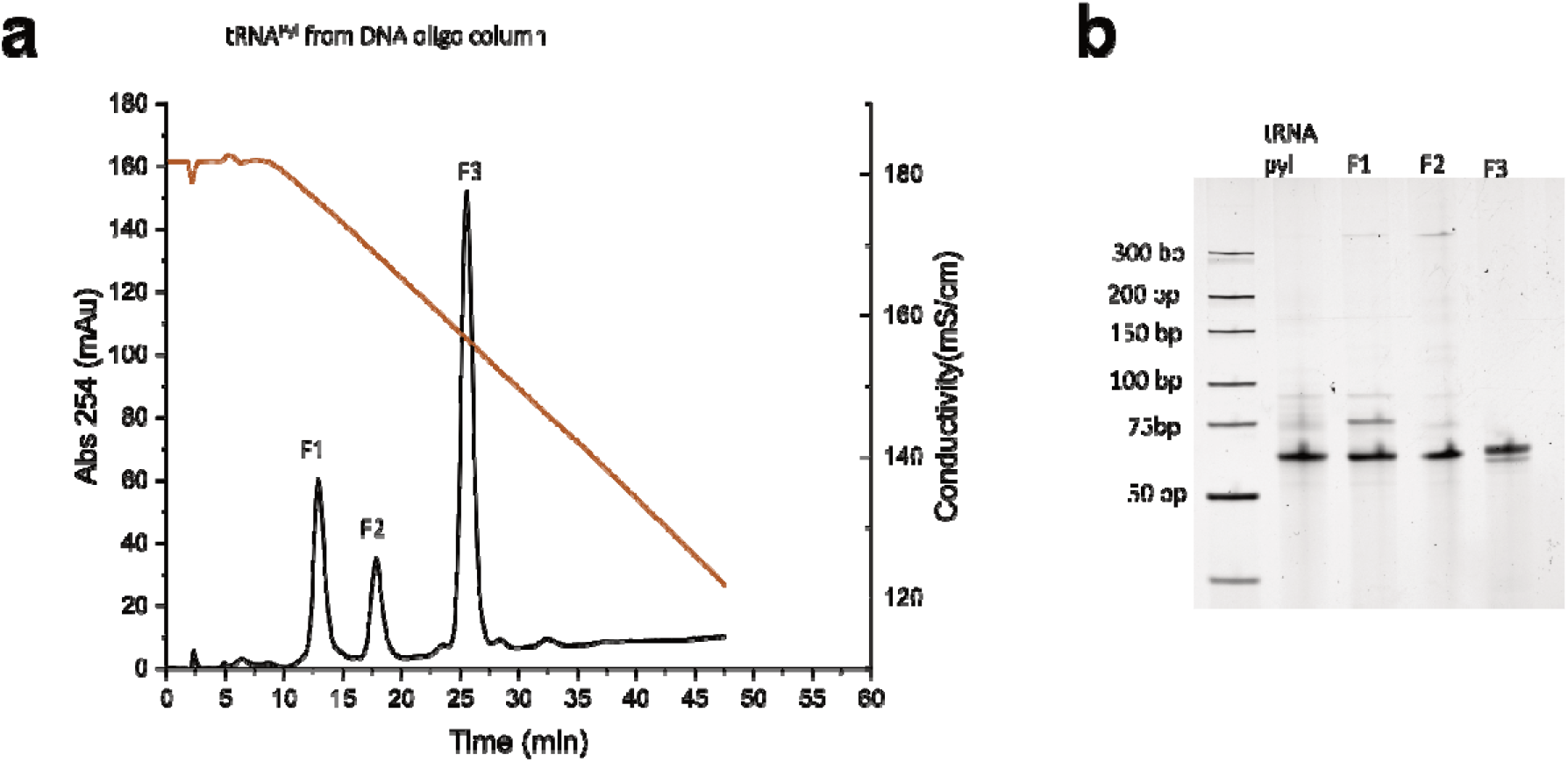
HIC separates fractions of purified *in vivo Ma*-. **(a)** Hydrophobic interaction chromatography (HIC) chromatogram of DNA-column purified *in vivo Ma*-. Column purified *in vivo Ma*-separated out into the three peaks, labeled as F1-F3. **(b**) Urea-PAGE of separated *in vivo Ma*-species. Lane assignments from left to right. Lane 1: RNA ladder, Lane 2: DNA-column purified *in vivo Ma*-, Lane 3: Fraction 1 of HIC-purified *in vivo Ma*-, Lane 4: Fraction 2 of HIC-purified *in vivo Ma*-, Lane 5: Fraction 3 of HIC-purified *in vivo Ma*-.

**Figure S7.**
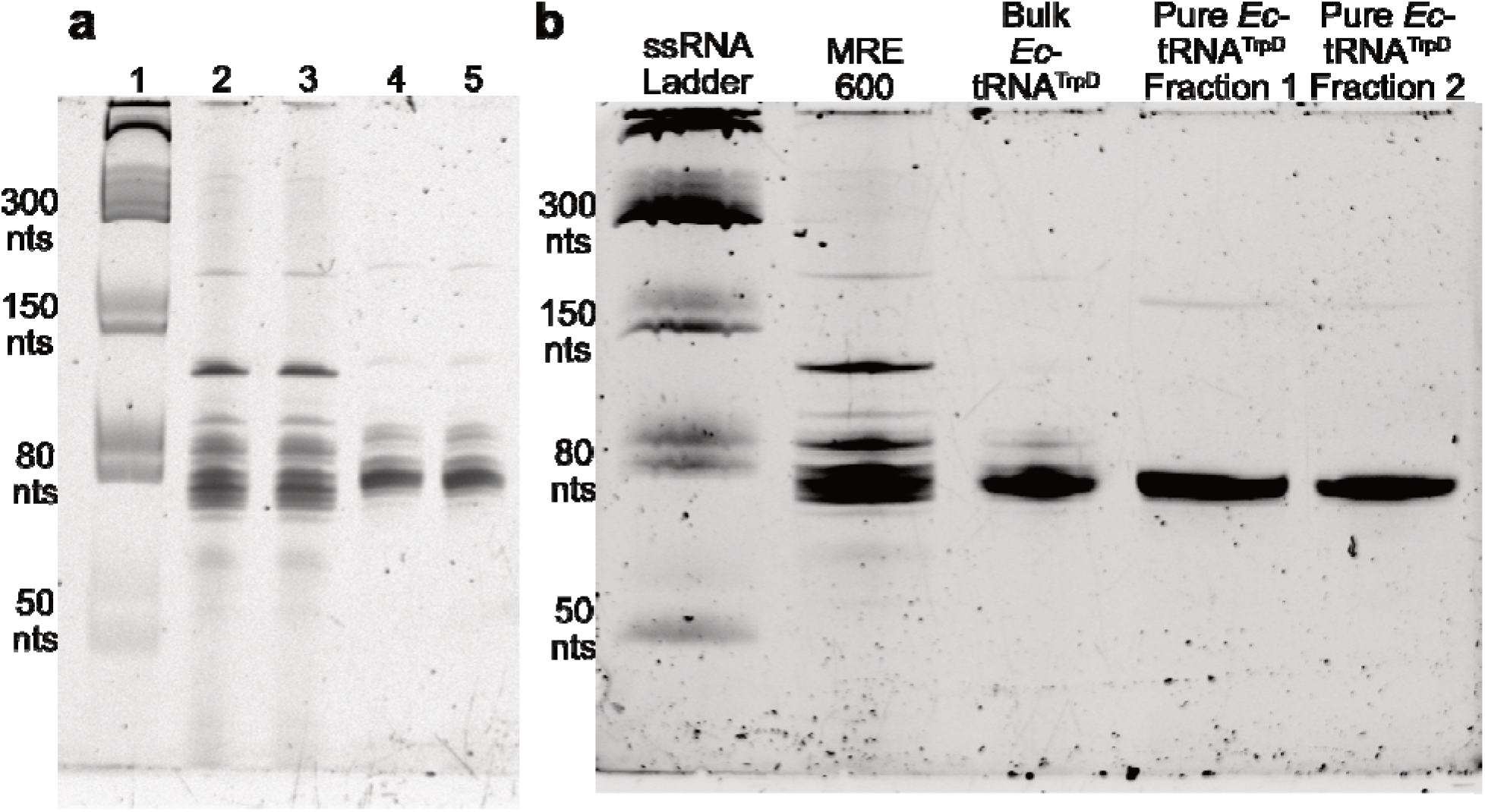
*Ec*-is greatly overexpressed and highly pure. **(a)** Urea-PAGE gel of overexpressed *Ec*-. Lane 1: ssRNA ladder, Lanes 2 and 3: MRE600, Lanes 4 and 5: overexpressed *Ec*-. **(b)** Urea-PAGE gel of purified *Ec*-. Lanes with purified *Ec*-contain either the first or second purification fractions.

